# Cell boundary confinement sets the size and position of the *E. coli* chromosome

**DOI:** 10.1101/348052

**Authors:** Fabai Wu, Pinaki Swain, Louis Kuijpers, Xuan Zheng, Kevin Felter, Margot Guurink, Debasish Chaudhuri, Bela Mulder, Cees Dekker

**Author notes:** Correspondence should be addressed to Cees Dekker and Bela Mulder 18.

## Abstract

While the spatiotemporal structure of the genome is crucial to its biological function, many basic questions remain unanswered on the morphology and segregation of chromosomes. Here, we experimentally show in *Escherichia coli* that spatial confinement plays a dominant role in determining both the chromosome size and position. In non-dividing cells with lengths up to 10 times normal, single chromosomes are observed to expand more than 4 fold in size, an effect only modestly influenced by deletions of various nucleoid-associated proteins. Chromosomes show pronounced internal dynamics but exhibit a robust positioning where single nucleoids reside strictly at mid-cell, while two nucleoids self-organize at ¼ and ¾ cell positions. Molecular dynamics simulations of model chromosomes recapitulate these phenomena and indicate that these observations can be attributed to depletion effects induced by cytosolic crowders. These findings highlight boundary confinement as a key causal factor that needs to be considered for understanding chromosome organization.

## Introduction

Chromosomes are spatially confined by physical boundaries. While interphase eukaryotic chromosomes reside in distinct territories within the nucleus (Bolzer et al., 2005), bacterial nucleoids occupy a large subvolume of the cytoplasm that is itself bounded by the cell membrane (Kellenberger et al., 1958). Historically, boundary confinement had been considered to be the sole factor constraining the structure of the bacterial and interphase-eukaryotic chromosomes, in contrast to the intrinsically condensed rod-shape eukaryotic chromosomes in metaphase. Studies in the past few decades revised this view by showing that chromosomes in all cells types and all phases of the cell cycle are structurally organized by various types of proteins interacting with DNA (Bickmore and van Steensel, 2013; Luijsterburg et al., 2006; Peeters et al., 2015). However, it remains elusive how the size of chromosomes is precisely determined in bacteria, archaea, and interphase-eukaryotic cells. Similarly, a general understanding of mechanisms underlying chromosome positioning in bacteria without mitotic spindles is lacking. This is largely due to the fact that to date the confinement-dependent effects could not be controlled independently, making it hard to disentangle the various proposed mechanisms.

The 4.6-Mbp circular chromosome of the rod-shaped *E. coli* is generally visualized as an ovoid nucleoid, occupying ~60% of the cell volume. PALM/STORM-type super-resolution microscopy was unable to resolve its detailed architecture (Wang et al., 2014) due to its small size and fast dynamics, whereas live cell imaging of a widened *E. coli* allowed an expansion of the ellipsoidal nucleoid into a torus that exhibited a strong density heterogeneity (Wu et al., 2018). This finding is consistent with various approaches indicating that *E. coli* chromosome organizes into a filamentous bundle with non-crosslinked left and right arms flanking the origin of replication, although the exact conformation of the arms can differ depending on nutrient conditions, cell width, and cell cycle (Fisher et al., 2013; Niki et al., 2000; Wang et al., 2006; Wiggins et al., 2010; Youngren et al., 2014). By contrast, some other bacteria such as *C. crescentus* show two arms that are crosslinked by condensin SMC protein complexes, but the individual arms are likely to also organize into filaments as inferred from 3C data (Umbarger et al., 2011). These studies of the shape and topology of bacterial chromosomes converge to a picture where in elongated bacterial cells, an internally compacted chromosome, with or without arm crosslinking, is constrained by the lateral cell wall into an ellipsoidal shape. Many proteins have been found to be associated with the internal compaction of DNA in bacteria, including nucleoid-associated proteins (NAPs, such as HU, Fis, and H-NS (Dame et al., 2006; Schneider et al., 1997; van Noort et al., 2004)) and structural maintenance of chromosomes proteins (SMCs, such as MukBEF in *E. coli* (Badrinarayanan et al., 2012; Lioy et al., 2018; Nolivos et al., 2016)). It however remains elusive how these proteins contribute to the overall size of the chromosome, even at the qualitative level.

The mechanism of chromosome positioning within the *E. coli* cell also remains an open question. During a cell cycle, a single nucleoid localizes around the cell center before DNA replication, while sister chromosomes localize to the two cell halves after they are replicated and segregated (Niki et al., 2000). So far, three main classes of mechanisms have been considered in the positional homeostasis and sister segregation of *E. coli* chromosomes: 1) physical effects of the intrinsic DNA polymer conformation and mechanics, 2) external forces acting on the whole chromosome, and 3) external forces acting on the OriC-proximal region. Numerical simulations showed that two long polymers can spontaneously separate from each other due to conformational entropy (Jun and Mulder, 2006), whereas dynamic imaging led to a proposal that chromosomes in live cells might be mechanically strained and repulse each other like loaded springs (Fisher et al., 2013). Other models proposed transertion (the tethering of DNA to the membrane through transcription-translation-coupling of transmembrane proteins (Woldringh, 2002)) and a coupling to the Min system (binding of DNA by membrane-bound MinD proteins which oscillate between the two poles (Di Ventura et al., 2013)). Finally, the Ori region is the first to be replicated and segregated during the cell cycle and it showed distinct localization patterns (Kuwada et al., 2013; Niki et al., 2000), prompting hypotheses that chromosome segregation and positioning are dictated by mechanisms acting on or near Ori. Various factors were proposed to drive Ori migration, although both the potential binding sites and the potential force-generating mechanisms still remain to be further elucidated (Kuwada et al., 2013; Nolivos et al., 2016; Yamaichi and Niki, 2004). Broadly speaking, it remains unclear whether chromosome segregation and positioning primarily rely on intrinsic or extrinsic driving forces, and whether these forces act locally or globally.

The study presented here is inspired by the increasing realization that the behavior of cellular structures is governed not only by specific molecular interactions, but also by the generically aspecific physical properties of the intracellular environment such as molecular crowding (de Vries, 2010; Ellis, 2001; Pelletier et al., 2012; Zhou et al., 2008) and by the boundary geometry (Young, 2006). In particular, mechanisms involved in cell growth and division depend on cell geometry to achieve organizational homeostasis (Hussain et al., 2018; Minc et al., 2011; Wu et al., 2015b). Given the fact that the chromosome occupies a large fraction of the total cell volume, it stands to reason that chromosome sizing and positioning should be understood in the context of cell size and cell shape.

Here, we study the size and position of a single nonreplicating chromosome in *E. coli* cells that range in length from 2 to 30 microns. We explore the principles by which chromosomes respond to cell size change and disentangle the roles of extrinsic and intrinsic factors to elucidate the underlying physical mechanism. We first combine genetic perturbation and quantitative imaging to show that the *E. coli* chromosome can reach a significantly larger size that depends nonlinearly on cell length, even though it is not in direct physical contact with the cell poles. Various nucleoid-associated proteins are shown to play secondary roles in quantitatively modulating the nucleoid-cell length relation. We use molecular dynamic simulations to show that depletion forces arising from molecular crowding provide a plausible mechanistic basis for capturing this behavior. We next investigate the morphological and positional dynamics of chromosome at various length scales. We find that in all cell lengths, a single nucleoid is positioned precisely at the cell center, whereas two sister chromosomes are positioned, non-self-evidently, at the ¼ and ¾ locations along the cell length. This persistent chromosome positioning is independent of Ori localization and of other proposed membrane-associated mechanisms, and can be recaptured by simulations, which identify the intrinsically slow global diffusion of the chromosomes and the entropically favorable distribution of newly synthesized crowders as the governing factors.

## Results

### Maintaining a single chromosome in a growing cell allows studying the effects of boundary confinement

In *E. coli* cells at steady-state growth conditions, the DNA replication is tightly regulated to scale the DNA copy number with the cell volume (Si et al., 2017), making it hard to probe the effect of cell-size changes on the size of a single chromosome. Here, we decouple DNA replication and cell growth so as to obtain cells that maintain only a single chromosome copy while sustaining a continued growth to very long lengths. Using a *dnaC2(ts)* mutant (Saifi and Ferat, 2012), a rapid shift from a permissive (30°C) to non-permissive temperature (40°C) will disable DnaC’s function in loading DnaB, an essential component of the replisome, which in turn prevents the cell from initiating new rounds of DNA replication. A second element of our approach is that we prevent cell division at any stage of the growth by adding cephalexin, an antibiotic which inhibits enzymes responsible for the septum cell-wall constriction. The nucleoids in the cells were labeled by HU-mYPet, which are endogenously expressed fluorescent-fusion proteins of a NAP that binds DNA in a sequence-nonspecific manner (Wery et al., 2001; Wu et al., 2015a). Origin- and terminus- proximal foci were labeled by fluorescent repressor-operator systems (FROS), as described previously (Reyes-Lamothe et al., 2008).

We inoculated these bacteria in microfabricated channels (Wu et al., 2015b) that were 1-μm wide, 1-μm high, and 60-μm long (Fig. 1A, see Methods). These channels guided single *E. coli* cells to grow linearly in one dimension. As cell division was prevented, cells containing a single chromosome could reach very large lengths of 20-30 μm. Supplementing the agarose pad with chloramphenicol to inhibit translation led to immediate cell growth arrest (Fig. S1A, S1B), in line with the recent finding that functional accumulation of cell mass underlies cell growth even when DNA replication initiation is inhibited (Si et al., 2017). These single-nucleoid *dna2(ts)* cells form the core system for studying the effects of boundary confinement on the bacterial chromosome.

**Figure 1.**
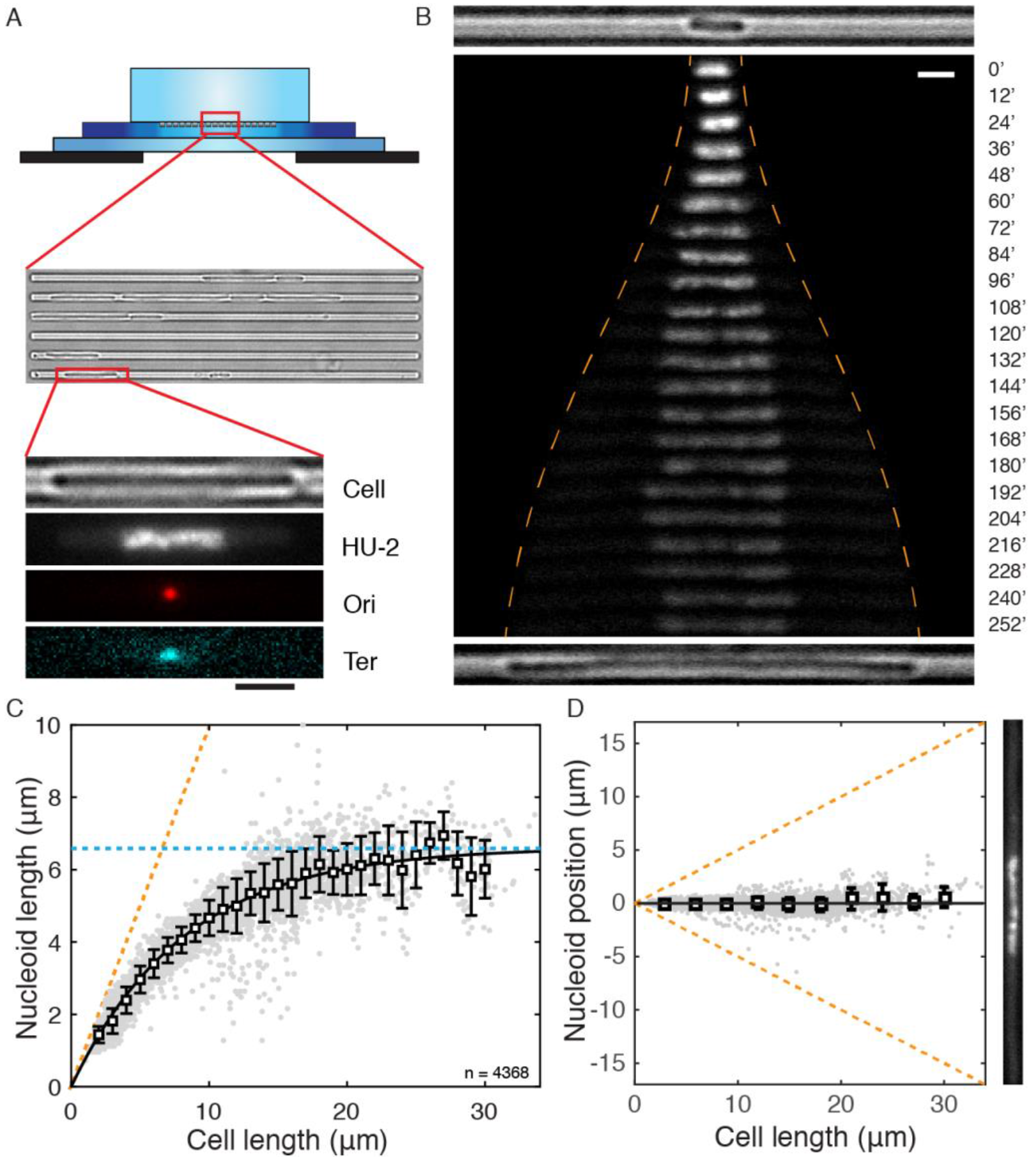
Chromosome size and positioning are dependent on cell size in *E. coli.* A. Schematic of the experimental set-up. Top illustration shows the cross section of the device composed of an agarose containing nutrient and drugs (top), a thin PDMS layer containing 1-μm-wide channels containing *E. coli* bacteria (middle), and a glass coverslip (bottom). Bottom panel shows, from top to bottom, a cell, its nucleoid, and the Ori and Ter loci, respectively. B. Time-lapse images of a HU-mYPet labeled chromosome as it expanding with cell growth at nonpermissive temperature defected in DNA replication initiation. The orange dash line indicate cell the positions of the cell poles. Time is indicated in minutes. Top and bottom panels respectively show the bright-field images of the cell at t=0’ and t=252’. C. The length of single nucleoids in relation to the cell length. Grey dots are single data points (n=4585). Squares and error bars are mean and standard deviations calculated with a bin size of 1 μm. Line shows an exponential decay (decreasing form) fit of the mean values *L*_*nucieoid*_ = 6.61*(1-exp(−0.12\**L*_*cell*_)). Orange dash line denotes a scenario where nucleoid occupies full cell length. Blue dash line indicates the maximal (intrinsic) cell length of 6.6 μm. D. Localization of nucleoid center of mass in relation to cell center. Squares and error bars are mean and s.d. values calculated with a bin of 1 μm, plotted every 3 μm. n = 4585. An image of a nucleoid in a long cell is exemplified at the right. Scale bars in A and B, 2 μm

### Nucleoid size scales nonlinearly with cell size

Systematic manipulation of the cell size allowed measuring the response of the nucleoid length to the degree of longitudinal confinement by the cellular boundary. Shown in Fig. 1B, a 2.8-μm-long cell at inoculation contains a single 1.6-μm-long nucleoid. As cell growth became apparent, the nucleoid did not retain this size, but instead started expanding longitudinally. The initial phase of nucleoid expansion was pronounced, doubling in length in an hour as the cell length doubled, indicating a near linear relation. In the following time course of cell growth, however, the chromosome expanded even further in a nonlinear way, ultimately reaching a length of 6.6 μm, about 4 times larger than its initial length. While the cell size and nucleoid size increased, the total number of nucleoid-bound HU-mYPet is steadily maintained (Fig. S1C), resulting in a drop of HU-mYPet intensity on the expanded nucleoid as well as a concomitant increase of it in the cytosol (Fig. 1B). The dramatic nucleoid-size expansion was surprising, as it was not predicted by the existing body of literature attributing chromosome size of bacteria to a combined effect of protein-mediated intra-nucleoid interactions (Lioy et al., 2018) and extrinsic cytosolic crowding (Pelletier et al., 2012), and thus warrants a thorough quantitative and mechanistic investigation.

We quantified the nucleoid-cell length relation in 4585 single-cell snapshots collected at different stages of cell growth. This led to a nucleoid-cell length relation that is well described by an exponential approach to saturation at 6.6 ± 0.2 μm, i.e. *L*_*nucleoid*_ = *L*_*max*_ *(1 - e^−Lcell/Lc^)* (Fig. 1C, coefficient of determination R^2^=0.97, Lmax = 6.6 ± 0.2 μm, Lc= 8.3 ± 0.5 μm, errors show 95% confidence). This fit captured both the early stage of near-linear increase of nucleoid size with cell size as well as the slowing down of expansion as cells grew larger until it approached saturation when the cells reached a length above 17 μm. This saturating behavior indicates that the nucleoid has an intrinsic length of 6.6-μm in the cylindrical cell geometry in the absence of longitudinal confinement.

### The nucleoid localizes strictly at mid cell position

Single nucleoids were found to strictly localize at the mid-cell position with a striking accuracy. As shown in Fig. 1D, the nucleoid center of mass is observed to coincide with the cell center, on average deviating from the mid-cell position over a distance less than 4% of the cell length (Fig. 1D). It is to be noted that, in conjunction with the above-described nonlinear relation between nucleoid and cell length, a very significant nucleoid-free cytosolic volume is observed near the two cell poles, whose size increased continuously without any saturation with cell length (Fig. S1D). This poses an intriguing question on how the nucleoid appears to “sense” the polar cell walls without any direct physical contact, a sensing that appears effective over long distances and remains operative beyond the cell length range within which the nucleoid length changes.

### The nucleoid contracts in size upon cell division

Given that a wide range of proteins was previously proposed to bind to DNA and influence the DNA compaction at various levels, it is conceivable that their concentrations or activities can quantitatively affect the chromosome size under the altered DNA/cytosol content ratio in our experiments. If confinement alone, rather than any potential changes in the activities of DNA-binding proteins or the overall degree of molecular crowding in the cytosol, were to determine the quantitative response of the nucleoid size to cell size observed above, the nucleoid would be expected to contract when the cell size were to be reduced.

To verify this experimentally, we examined the nucleoid sizes before and after cell division in a *ΔslmA/dnaC2* mutant at different times (Fig. 2 and Fig. S2). SlmA is known to bind DNA and depolymerize FtsZ to prevent cell division at positions across the nucleoid (Bernhardt and de Boer, 2005). When SlmA is omitted in our single-nucleoid cells (in the absence of cephalexin), the cells were found to frequently divide at the nonpermissive temperature (Fig. S2A), and, interestingly, they were observed to distribute DNA copies unequally among progenies. Notably, only the daughter cells that inherited DNA continued to grow. The *ΔslmA/dnaC2* mutant thus demonstrated that the single-nucleoid cells are metabolically active as that cell growth is fueled by active transcription from DNA.

**Figure 2.**
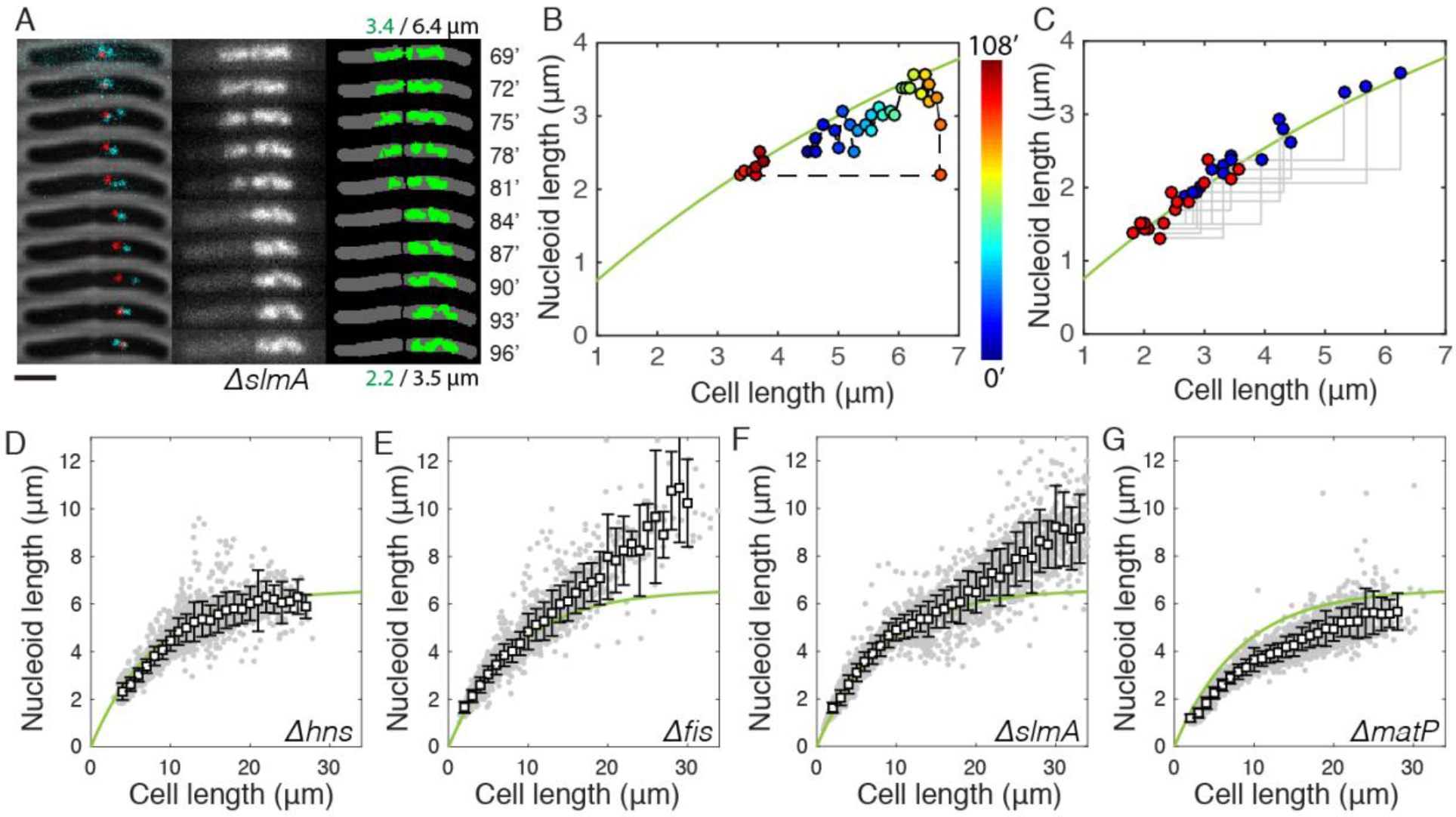
Cell-size-dependent chromosome sizing under extrinsic and intrinsic perturbations. A. Time-lapse images *slmA/dnaC2* cell division at 3-minute intervals. Left: Ori (red) and Ter (cyan) foci overlayed on phase contrast images. Middle: DNA visualized through HU-mYPet. Right: Binary overlay of cell body and the nucleoid. Numbers at top and bottom indicate nucleoid/cell lengths in the first and last frame. B. Nucleoid length versus cell length for the cell shown in A during a full growth and division cycle. Color bar shows time. The green line is identical to the dependence in Fig. 1C. Nucleoid length versus cell length before (blue) and after (red) cell division in *ΔslmA/dnaC2* cells (n=16). The green line is identical to the dependence in Fig. 1C. D-G. Nucleoid length versus cell length in cells respectively lacking *hns* (n=2175), *fis* (n=2291), *slmA* (n=3125), or *matP* (n=2678) genes. The smooth green line represents the wildtype data as shown in Fig. 1C, for comparison.

These above manipulation thus led to a ‘reverse’ control system for examining chromosome sizing upon cell *shortening*, as the nucleoid traversed from one long cell into one shorter daughter cell. Time-lapse imaging at 3-minute intervals showed that the single genome copy residing in the mother cell was first pinched by the constricting septum and then rapidly translocated to one compartment before cell scission (Fig. 2A, more examples see Fig. S2B). These translocations are unidirectional (always towards the cell halves containing the Ori, Fig. S2C) and occurred with a 5kbp/s maximum speed (Fig. S2D), in agreement with the *in vitro* measured speed of DNA translocase FtsK (Saleh et al., 2004). Strikingly, the nucleoids became *smaller* in the (smaller) daughter cell, but again did not fill up the volume of the latter (Fig. 2A, S2A, S2B). Figure 2B shows the quantitative analyses of individual cell division events, which all yielded nucleoid-cell size data from mother-daughter pairs that collapse onto the same curve that describes the chromosome expansion with cell elongation (Fig. 1C). Notably, nucleoid contraction took place in a ~5-10-minute time frame near the septation event (Fig. 2A, S2B), too short for significant changes in the cellular crowding, metabolic state, or NAP concentrations to occur. Quantitative mapping of single nucleoid/cell size over time showed that they consistently fluctuate around the same curve (Fig. 2C), even in cells that underwent two consecutive growth-division cycles (Fig. S2E). Hence, we conclude that a change in longitudinal confinement alone is responsible for the observed rapid and reversible nonlinear scaling of the nucleoid size with cell size.

### NAPs exhibit modest effects on the nucleoid size

Next, we explored the roles of intrinsic packaging agents on the nucleoid size by independently omitting various NAPs in our wildtype strain background described in Fig. 1. Specifically, we probed the abundant and well-studied NAPs Fis and H-NS, which distribute across the genome and have long been proposed to induce chromosome compaction (Dame et al., 2006; Schneider et al., 1997), as well as SlmA and MatP, which target binding sites away from and close to the terminus region, respectively (Bernhardt and de Boer, 2005; Mercier et al., 2008).

Nucleoids of the *Δhns cells* exhibited a nonlinear increase with cell size (Fig. 2D) that, remarkably, was almost identical to *NAPs*+ cells (*NAPs*+ denotes the control strain described in Fig. 1), showing a saturation at 6.7 ± 0.2 μm (R^2^ = 0.98). This finding is unexpected as H-NS has long been thought to play an essential role in chromosome compaction and was recently observed to promote short-range interactions. Through PCR and sequencing, we found no extra copy of *hns* gene elsewhere in the genome and no mutation in the hns-paralog *stpA* gene. We also examined the physiological effect of *Δhns* and found that, at the permissive temperature of 30°C, these cells grew much more slowly than *hns+* cells (doubling time 165 vs. 83 minutes). We thus conclude that H-NS proteins, despite being essential for the homeostasis of cellular metabolism as a global transcription repressor, have virtually no effect on the global nucleoid size.

Omitting Fis and SlmA also showed little effect in cells shorter than 15 μm but removal of either of these NAPs was observed to lead to clearly longer nucleoids compared to *NAPs*+ strains in cells longer than 15 μm, see Fig. 2E and 2F. At the maximum cell length of ~30 μm, the nucleoid length reached 10.2 ± 1.8 μm and 9.2 ± 1.7 μm, respectively, significantly above the 6.6 μm plateau for wildtype nucleoids. These data strongly indicate that Fis and SlmA both play a role in determining the degree of intrinsic DNA-cross-linking that contribute to the observed maximal nucleoid length of 6.6 μm. The effect of Fis can be attributed to its previously reported functions of bending DNA *in vitro* (Pan et al., 1996) and stabilizing supercoils *in vivo* (Schneider et al., 1997). The effect of SlmA is surprising as its role in chromosome organization was so far barely investigated, although 3C data did show that SlmA-binding sites have higher interactions with their neighboring sequences (Cagliero et al., 2013). Despite the strong effect at larger cell lengths, however, in cells with a size smaller than 15 μm (5 times the regular cell sizes), the strong effect of boundary confinement overruled any effects of changes in local DNA crosslinking by Fis and SlmA.

Omitting MatP led to a 20% reduction in nucleoid size compared to wildtype (Fig. 2G). This observation is in line with recent finding that MatP proteins modulate the actions of MukBEF (Lioy et al., 2018; Nolivos et al., 2016) and are responsible for inducing a thin Ter region (Wu et al., 2018), rather than condensing the Ter region (Dupaigne et al., 2012; Mercier et al., 2008). Unlike Fis and SlmA, the effect of MatP is apparent across all cell lengths, showing that its role in condensing the chromosome acts in parallel to the effect of boundary-confinement and is relevant to the nucleoid size in regular cells at steady-state growth conditions.

### Polymer modeling captures the sizing and positioning of nucleoids when including molecular crowders

To explore the physical mechanisms underlying the experimentally observed intrinsic nucleoid length, i.e. the 6.6-μm saturation, as well as its compaction by longitudinal confinement, we carried out molecular dynamic simulations based on a simple polymeric chromosome model (Chaudhuri and Mulder, 2012; Jun and Mulder, 2006; Reiss et al., 2011)). This model captures a loop-based chromosomal organization principle (Ganji et al., 2018; Postow et al., 2004) by considering a self-avoiding polymer consisting of a circular backbone chain to which a large number of side-loops are attached (Fig. 3A, Fig. S3A), a so-called “bottle brush” structure (Rathgeber et al., 2005). The impact of the side-loops is further coarse-grained by representing their contribution in terms of an effective repulsive Gaussian core interaction (Stillinger, 1976) between the backbone monomers (Fig. 3A) (see Methods section for model details). The model partitions the 4.6-Mbp genome into a circular main chain to which ~ 600 loops are attached at uniform separation and of equal size close to the experimentally reported mean loop size (Postow et al., 2004). We simulated such polymers inside a cylindrical volume of 1.0 μm diameter and variable lengths, with different concentrations of crowder molecules.

**Figure 3.**
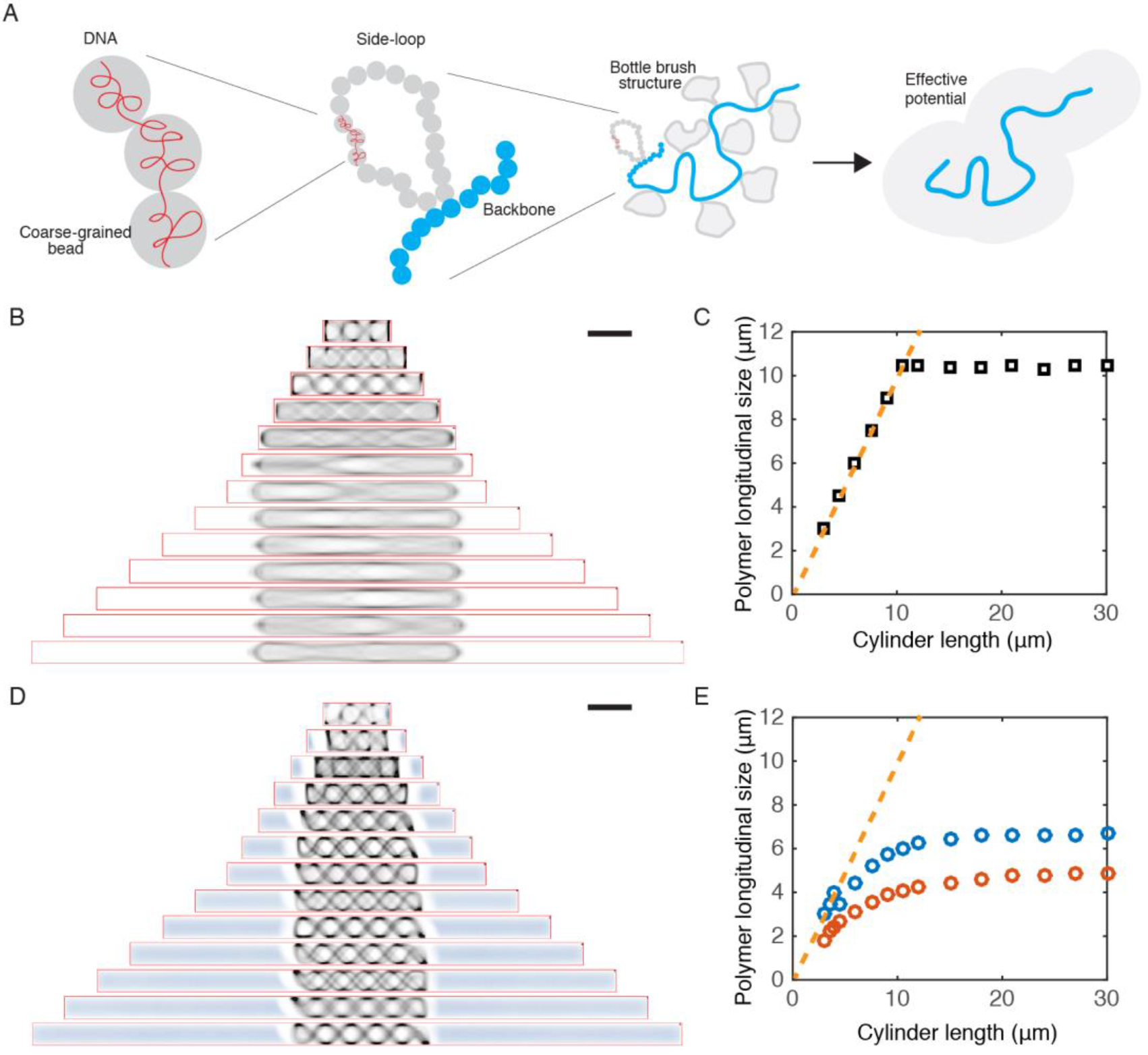
A polymer model captures the effect of boundary confinement on nucleoid size and position. A. Schematic of the construction of our coarse-grained polymer model of bottle-brush type, with a bead chain circular backbone and side loops represented by a parametrized effective potential. B. Time-averaged conformations of our model chromosome simulated in cylindrical cells of different lengths in the absence of depletants. C. Longitudinal size of the modeled chromosome polymer as a function of cell size, simulated without depletants. Note that the Gaussian core size is fixed in the simulations. Dashed orange line indicates the cell length. D. Time-averaged conformations of our model chromosome simulated in cylindrical cells of different lengths in the presence of depletants at density of 212 μm^−3^. E. Longitudinal size of the modeled chromosome polymer as a function of cell size, simulated with two different concentrations of depletants. Note that the Gaussian core size is fixed in the simulations. Blue indicates a depletant density of 212 μm^−3^, and red indicates a depletant density of 1060 μm^−3^. Dashed orange line indicates the cell length. Scale bars in B and D, 2 μm.

Our numerical simulations suggest that cell-size sensing by chromosomes can arise through its interactions with cytosolic crowders (Fig. 3B-E). We first carried out simulations without cytosolic crowders, as was done in all previous modelling work on bacterial chromosomes (Chaudhuri and Mulder, 2012; Jun and Mulder, 2006; Wiggins et al., 2010). We observed that the polymer pushed against the poles of the cylinder and formed helical conformations, until the cylinders were sufficiently long to allow the polymer backbone to completely stretch out (Fig. 3B, 3C). This is notably different from the experimental observations. Next, we incorporated depletions effect from cytosolic crowders by adding so-called non-additive crowder particles (Dickinson 1979, Dijkstra et al., 1998). Shown in Fig. 3D and 3E, the crowders spontaneously segregate from the DNA polymer spatially and localize to the peripheries of the confining cell.

Upon elongating the cell, we observe two key effects of the crowders on the longitudinal size of the chromosome. First, crowders that were introduced exert an inward pressure on the chromosome generating a much more compact shape, as well as a central localization (Fig. 3D). At the local scale, the backbone was observed to buckle at many locations along the polymer (Fig. S3A), effectively reducing the length of the backbone when observed at lower resolution. At the global scale, the backbone showed a helical morphology with micron sized helical pitch even in the longest cylinders (Fig. 3D), unlike in simulations without crowders, where the backbone entirely stretched out (Fig. 3B). Interestingly, such a helical conformation was also captured by our structured illumination microscopy (SIM) images (Fig. S3B). Secondly, the simulation estimate of chromosome size as a function of cell size was nonlinear and much more gradual, in much better agreement with experimental findings (Figure 3D). Numerically, the two simulation data sets shown in Fig. 3E yielded saturation values of 6.7 and 4.9 μm for two crowder densities, close to the experimentally observed value, which is gratifying in view of the simplicity of the model. While our elementary model with a uniform loop size and constant crowder density thus captures the experimentally observed trends remarkably well, further modeling of the profile of the nucleoid-cell length relation will benefit from refinements by additional factors including the heterogeneous and dynamic nature of both the DNA loop distribution (Fisher et al., 2013; Postow et al., 2004; van Loenhout et al., 2012; Wu et al., 2018) and the cytosolic particle sizes (Parry et al., 2014).

### Chromosome are strongly dynamic internally but weakly diffusive globally

In live cells, chromosomes exhibit strong intrinsic morphological dynamics. Time-lapse SIM imaging in live cells revealed rapid morphological transformations and density drifts within the long helical chromosomes at sub-minute time scale (Fig. 4A). The coefficient of variation (*C*_*v*_=*s.d./mean*) of the nucleoid length stayed rather constant at around *C_v_* ~ 0.13 across all cell lengths (Fig. 4B).

**Figure 4.**
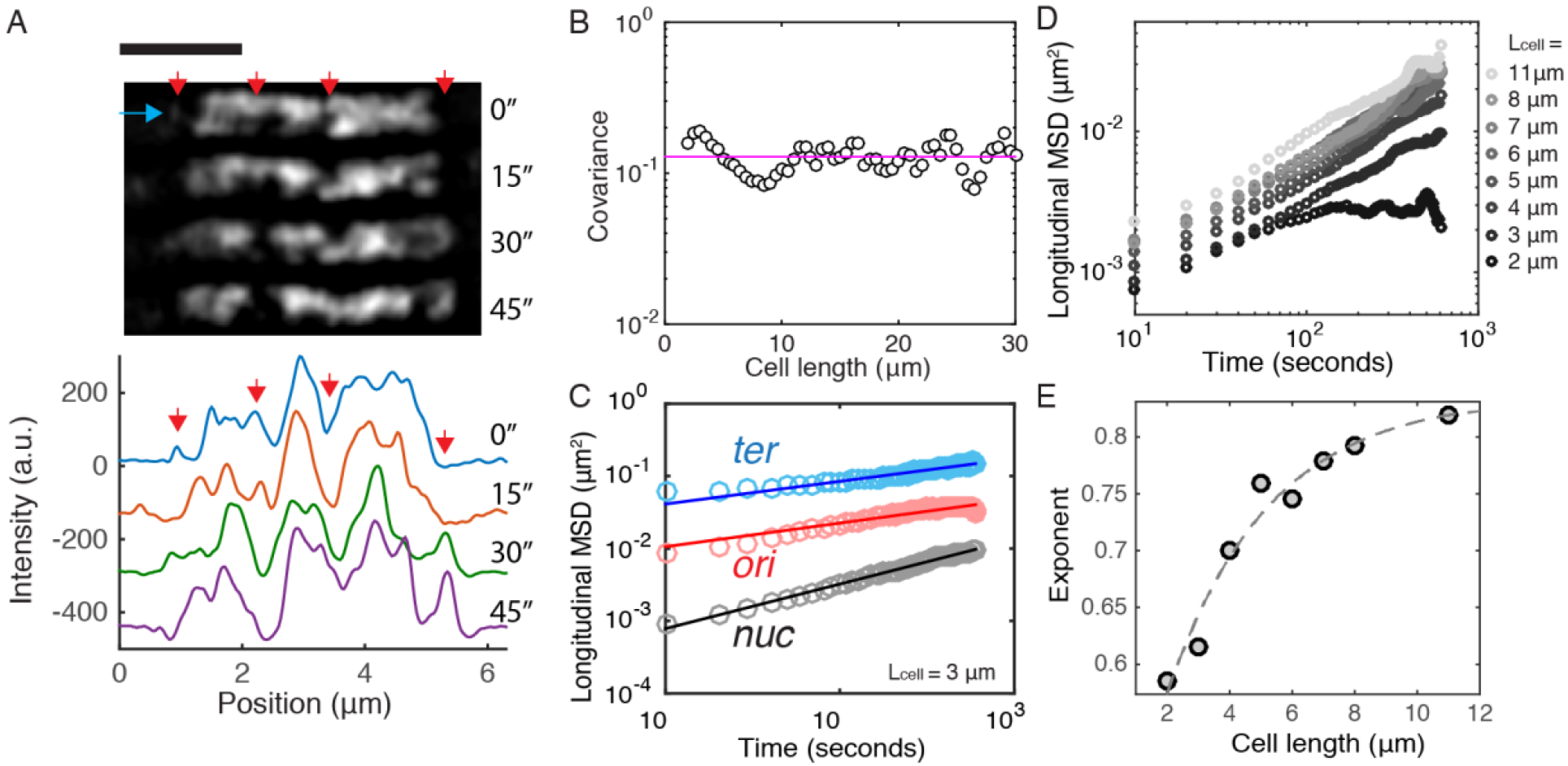
The *E. coli* chromosome is strongly dynamic internally but weakly diffusive globally. A. Structured Illumination Microscopy (SIM) images showing rapid density drifts and morphological changes of within a long nucleoid. The red arrows indicate areas with significant changes. The blue arrow indicates the cross-section along which intensity profiles are taken as displayed in the plot below the images. Scale bar, 2 μm. B. Comparison of the covariance of the nucleoid lengths in different cell lengths. The mean values is shown in magenta. C. Mean square displacement (MSD) of nucleoid center of mass (black), Ori foci (red) and Ter foci (cyan) along the long axis in 3-μm-long cells versus time. Circles indicate experimental data and lines indicate fits for subdiffusion. D. MSD of nucleoid center of mass versus time in different cell lengths. E. Exponent of the fits describing sub-diffusion of nucleoids in different cell lengths (Diffusion coefficients are all 1.9×10^−4^ μm^2^/s^α^). The dashed line denotes an exponential approach to saturation fit, *f(x)* = *0.84 - 0.48e*^*−0.31x*^. Note that for 2-μm cells the exponent was calculated for the first minute, where the profile follows the power law, before the trajectory plateaus.

We next compare the local and global behavior of the chromosome by measuring the mean-square displacement (MSD) in time lapse experiments for the Ori and Ter foci as well as for the chromosome center of mass (COM) at 10-second time resolution at 40°C. Figure 4C shows the data for 3-micron-long cells. The MSD of the Ori and Ter foci is seen to scale as a power law with time, as expected for sub-diffusion, <Δ*x*^2^> = *D t*^α^ where *D* is a generalized diffusion coefficient. The Ori and Ter traces are fitted by very similar exponents α (0.31 vs. 0.33, respectively), but they differ strongly in the diffusion coefficient D which is seen to be much larger (2×10^−2^ μm^2^/s^α^) for Ter than for Ori (5×10^−3^ μm^2^/s^α^). Interestingly, the COM of the entire nucleoid also followed a subdiffusive behavior, albeit with a much lower diffusion coefficient of 1.9×10^−4^ μm^2^/s^α^ and a larger exponent of 0.62. These data show that the diffusive behavior of the chromosome as a whole is distinct from its local dynamics. While, Local DNA loops are strongly dynamic, they are restricted to a certain region due to the polymeric nature of the chromosome as well as the local compaction density. By contrast, the chromosome is in principle free to explore the whole cellular space, but its large size and the high cytosolic viscosity together constrain its diffusivity.

We next examine how the longitudinal boundary confinement plays a role in the diffusivity of the chromosomes. It is commonly known that confinement affects the MSD due to the finite length that can be travelled. This is indeed observed in the shortest, 2-μm-long cells, where MSD saturates after 1 minute of imaging (Fig. 4D). In cells longer than 3μm, no saturation in MSD was observed within the 10 minutes duration of the experiments (Fig. 4D). Surprisingly, however, we observe an additional effect of confinement on the sub-diffusion behavior of the nucleoid COM: While it maintained a near-constant diffusion coefficient, it exhibited a pronounced dependence of the exponent that increased from <0.6 to >0.8 with increasing cell length (Fig. 4E).

### Persistent chromosome central positioning independent of Ori/Ter localization

The above data on chromosome dynamics suggests that while strong morphological dynamics of chromosomes can arise through active transcription and metabolism (Fig. 4A-B), confinement and crowding still have strong effect in constraining their global dynamics to sub-diffusion (Fig. 4C-E), contributing to their persistent positioning at long term (Fig. 1D).

Previous work suggested various Ori- and Ter-associated mechanisms to play a role in chromosome segregation and distribution (Danilova et al., 2007; Espéli et al., 2012). We thus analyzed the localization patterns of Ori/Ter loci positioning in our experiments during cell growth and compare that to nucleoid COM. Shown in Fig. 5A, Ori loci localize near the center of the cell, with standard deviation close to that of the nucleoid COM, whereas the localization of Ter loci in average are farther from the cell center. Quantitative analyses of fluorescent Ori loci revealed an accurate localization of the origin of replication to the nucleoid center in *wildtype* cells whereas Ter loci exhibited a larger spatial freedom (Fig. 5B and 5C).

**Figure 5.**
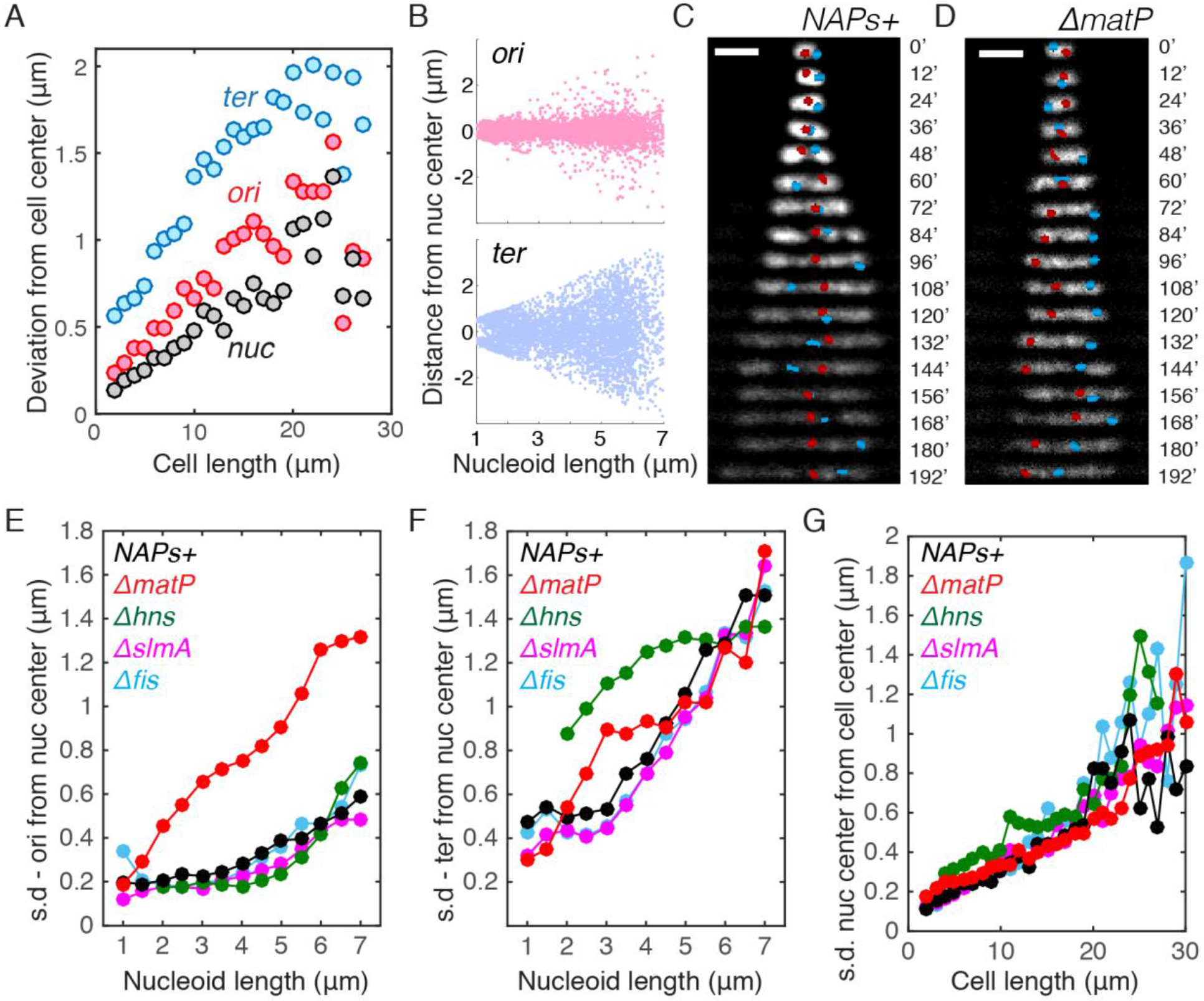
Persistent positioning of single chromosome independent of NAP-modulated substructuring. A. Deviation (mean square root distance) of the nucleoid center, Ori locus and Ter locus from the cell center in cells of different lengths. B. Distances of Ori / Ter loci from the center of nucleoids in relation to nucleoid length. C. Time-lapse images showing the positioning of Ori locus (red) and Ter (blue) locus in single nucleoids over time. Scale bars are 2 μm. D. Time-lapse images showing the positioning of Ori locus and Ter locus in single nucleoids over time for the ΔmatP strain. E. Deviation (mean square root distance) of the Ori foci from the nucleoid center in different mutants in different cell lengths. *NAPs*+ denote the control strain with all NAPs present. F. Deviation (mean square root distance) of the Ter foci from the nucleoid center in different mutants in different cell lengths. G. Deviation (mean square root distance) of the nucleoid center of mass from the cell center in different mutants in different cell lengths.

The above data suggest that nucleoid COM more accurately localize to the cell center than the labeled Ori locus. However, given that chromosomes are significantly larger and inherently less diffusive than an individual OriC locus (Fig. 4A), the causal relation between the localization of Ori region and nucleoid COM to the cell center remains insufficiently resolved. To elucidate it further, we examined the nucleoid loci and COM positioning in various NAP mutants, and found that *ΔmatP* cells lost the central localization pattern of the Ori foci (Fig. 5D-F, S4). This is consistent with recent finding that MatP regulates MukBEF and TopoIV to modulate Ori organization (Nolivos et al., 2016), and affect their local DNA structure (Wu et al., 2018). Surprisingly, the persistent localization of the nucleoid COM to the cell center did not alter in *ΔmatP* cells (Fig. 5G). In addition, the nucleoid COM was also observed to persist at the cell center in *Δhns* cells where Ter loci resided at the side of the nucleoid, and in *Δfis* and *ΔslmA* cells where Ori/Ter localizations are similar to the NAP+ strain (Fig. 5D-G, S4). Hence, the persistence of single chromosome at cell center is found to be independent of the localization of Ori or Ter region.

### Sister chromosomes position at ¼ and ¾ of all cell lengths

Next, we examined cells containing two chromosomes. Here, we observed a highly specific positioning of the two nucleoids in the cells. Upon sustained cell growth, the two sister chromosomes separated and accurately localized to the two quarter positions along the long axis, that is, at ¼ and ¾ of the cell length (Fig. 6A). This is by no means trivial, as *a priori* one might expect them to be free to localize anywhere along the cell length, provided they do not overlap. Or perhaps, one might have anticipated that on average they would localize near 1/3 and 2/3 positions. However, a ¼ and ¾ positioning pattern was robustly seen for almost all cells with two completely replicated chromosomes and, strikingly, this persisted for all cell lengths (Fig. S5A).

**Figure 6.**
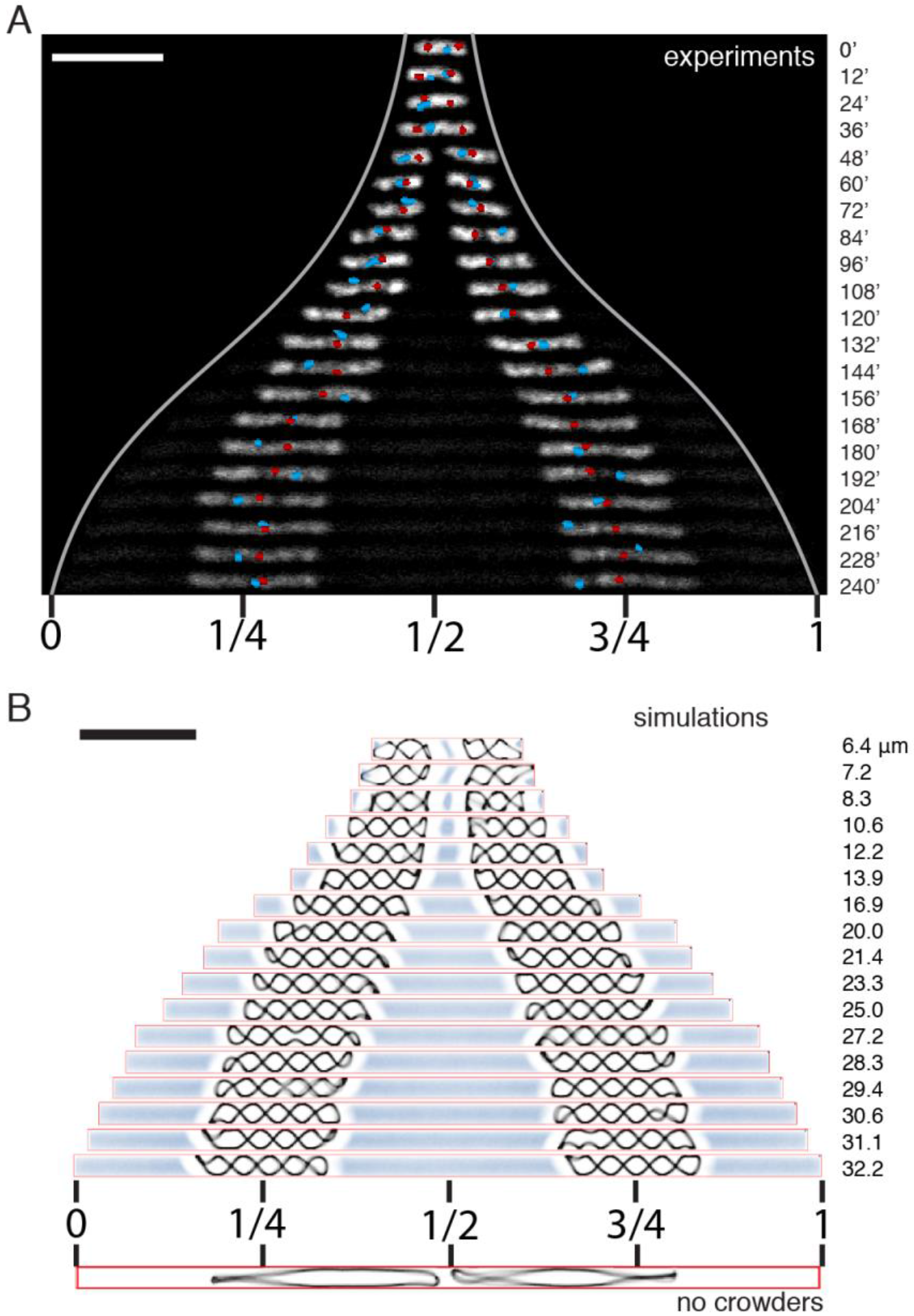
Positioning of two replicated chromosomes. A. Time-lapse images of nucleoid positioning in cells that contain two chromosomes. Cell poles are indicated by the light grew lines. Center and quarter positions in the final cell length is indicated below the image. Ori loci are shown in red, and the Ter loci are shown in cyan. B. 2D projection of simulated sister chromosomes that are moving apart due to cell growth and the associated depletant addition. Cell lengths are indicated at the right.
Scale bars, 5 μm.

The remarkable accuracy of the nucleoid localization prompted us to explore the possible role of active mechanisms that had been proposed. We first deleted the *minDE* genes in light of the proposal that Min oscillations may affect the positioning of chromosomes (Di Ventura et al., 2013). However, we found no effect (Fig. S5B). We next examined the involvement of transertion that might tether chromosomes to the membrane (Woldringh, 2002). To test this, we treated the elongated cells with a combination of chloramphenicol and rifampicin (see Methods) to inhibit both transcription and translation, but we did not observe change in nucleoid positioning (Fig. S5C, S5D). We conclude that these active mechanisms do not play a role in the nucleoid localization.

Subsequently, we explored the effect of entropic repulsion in sister chromosome segregation using molecular dynamics simulations of two copies of nucleoid in a growing cylindrical confinement (Fig. 6B, bottom). In absence of crowders, the chromosomes were initially able to localize to the ¼ and ¾ positions due to direct repulsion between the chromosomes in small cells, but proper spatial segregation failed for cells longer than 20 μm where the direct chromosomal overlap disappears beyond the length of two fully stretched nucleoids (grey lines in Fig. S5E). This approach thus did not fully recapitulate the experimental finding.

The correspondence to the experiments however greatly improved when we examined the effect of macromolecular crowding. Using Boltzmann-weighted insertion of new crowders (see Methods), we ensured that they were inserted homogeneously in the space outside of the chromosomes. As a result, the initial ¼ and ¾ positioning due to the direct repulsion between the chromosomes was maintained by a 1:2:1 partitioning of the crowders to the space between one cell end and the first chromosome, the space between the two chromosomes, and the space between the second chromosome and the other cell end. This resulted in a balanced compression force exerted on the chromosomes by the crowders and an effective repulsion between them, even in the longest cells beyond the regime of direct chromosomal overlap (Fig. 6B). Thus, a force generation due to entropic dispersion of the crowders promotes the ¼ and ¾ positioning for all cell lengths, including those beyond 20 μm, where the bare model without crowders failed (Fig. S5E).

The robust the ¼ and ¾ positioning is due to two partly history-dependent kinematic mechanisms, (i) direct inter-nucleoid repulsion in small cells, (ii) longer ranged effective repulsion between chromosomes through continued homogeneous protein production in the space outside of the chromosomes. Both of these driving mechanisms are entropic in origin.

## Discussion

In this paper we demonstrated how the size and position of *E. coli* chromosomes depend on the cell size. Quantitation and modeling of the chromosome-boundary relation allowed us to identify the driving forces that govern chromosome organization and disentangle the roles of diverse factors known to interact with DNA.

The first key finding of this study is that, without directly pushing against the cell poles, the *E. coli* nucleoid senses the level of longitudinal confinement and varies its size accordingly. This takes place in the case of chromosome expansion during cell growth, as well as in chromosome contraction at cell division. Our simple polymer model indicates that confinement acts on chromosomes by modulating the force balance induced by cytosolic crowders. Surprisingly, several NAPs that were previously found to induce DNA crosslinks were experimentally shown to play only secondary roles.

The extent to which the chromosome size reacts to changes in longitudinal confinement is surprising. The existence of a distinct nucleoid region within *E. coli* was reported as early as the 1950s (Kellenberger et al., 1958). As the nucleoid was seen to push against its cell envelope transversely, but not longitudinally, discussions on the effect of confinement primarily focused on how the small cell diameter influences the chromosome morphology (Fisher et al., 2013; Youngren et al., 2014), while the longitudinal compaction of nucleoid has been mainly considered to be determined by intrinsic packaging by NAPs and SMCs (Lioy et al., 2018). In principle, chromosome compaction can be well achieved by protein-mediated DNA-crosslinking alone (Luijsterburg et al., 2006). However, the merit of relying on confinement becomes apparent once we consider its physiological advantages. Strong protein-mediated DNA condensation can be found in metaphase eukaryotic cells or deep-stationary-phase bacterial cells, but such a highly packaged state imposes a disadvantage for its accessibility to transcription and replication machineries. However, by taking advantage of the confinement effect for physiologically relevant levels of crowding (de Vries, 2010; Ellis, 2001; Pelletier et al., 2012; Zhou et al., 2008), the chromosome can achieve a relatively small size with a modest level of intranucleoid organization while allowing both dynamics and accessibility.

Our quantitative data of the confinement effects in cells with various genetic perturbations have strong implications on the understanding of the intranucleoid interactions mediated by various NAPs. H-NS and Fis have been shown to bridge DNA and change its conformations *in vitro* (Dame et al., 2006; Schneider et al., 1997). Recent Hi-C studies also showed that they respectively promote short- and long-range DNA-DNA interactions (Lioy et al., 2018). The functional consequences of these interactions on nucleoid size were, however, not as expected. Here we showed that the interactions mediated by Fis and H-NS did not influence nucleoid size in cells smaller than 15 μm, which is 5 times larger than a regular G1-phase *E. coli* cell with a single nucleoid. This would suggest that the reported Fis- and H-NS-mediated DNA-DNA interactions are instead important in transcription regulation, in line with recent finding that Fis is essential for the emergence of transient domain boundaries across the dynamic genome in a live cell (Wu et al., 2018). A confinement-driven mechanism underlying nucleoid size homeostasis thus shows an advantage in tolerating changes in local DNA topology as influenced by transcription. The nucleoid size can, however, be tuned by MatP proteins, which expanded the nucleoid by 20% at all cell sizes. This can be explained by the recent finding that MatP reduces DNA compaction at Ter and Ori region (Wu et al., 2018). This study indicates that this structural modulation by MatP also appears to be essential for the internal conformation (Ori centering) of the nucleoid.

The second key finding of this study is that confinement-modulated depletion forces place the nucleoids persistently at a defined position. The depletion forces induced by cellular crowders, which are entropic in origin, appear weak enough to allow prominent morphological dynamics at the local scale, but strong enough to curb full-chromosome mobility at the larger scales. The essential role of depletion force that we observed is notably consistent with the recent prediction that a weak force, larger than purely entropic polymer-polymer repulsion force but much smaller than that generated by canonical motors, drives chromosome segregation in *E. coli* (Kuwada et al., 2013). It is also in line with recent experimental data showing that replicated chromosomes do not spatially segregate without cell growth (Woldringh et al., 2015). Clearly, the small magnitude of the force responsible for the positioning homeostasis of the chromosome allows it to be easily overcome by active ATP/GTP-driven processes that involve DNA transport across the cell length, such as FtsK-mediated DNA translocation ((Männik et al., 2017), also see Fig.2), or RecA-mediated DNA repair (Lesterlin et al., 2013). It is known that bacteria such as *C. crecentus* use active mitotic machineries to segregate chromosomes, raising the intriguing question whether mitotic/non-mitotic mechanisms result in different evolutionary advantages. We can speculate that whereas a motor-driven mechanism enables polar localization and daughter-cell differentiation, an entropy-driven mechanism is arguably more free-energy efficient.

All cellular processes occur in the context of confinement. Recent studies of the effect of boundary geometry largely focused on nonequilibrium self-organized systems such as reaction-diffusion patterns (Wu et al., 2015b) and molecular-motor-driven active fluids (Wu et al., 2017). Here we showed how the confinement determines the chromosome size, dynamics, as well as positioning. These findings have broad implications on the organization of bacterial, archaeal, and eukaryotic-interphase chromosomes under their confining envelopes, as well as the confinement-dependence of diffusivity in cytoplasm in general.

## Acknowledgement

We thank Erwin van Rijn, Jeremie Capoulade, Dimitri de Roos, Jelle van der Does, Alexandre Japaridze, Jakub Wiktor, Anne Meyer, and the staff at Kavli NanoLab for technical support and discussions. The work was supported by the Netherlands Organisation for Scientific Research (NWO), the NWO/OCW programs NanoFront and Basyc, and by the European Research Council Advanced Grant SynDiv (No. 669598). D.C.’s work was supported by SERB, India through grant number EMR/2016/001454.

## Author contributions

F.W. and C.D. designed the experiments. F.W., L.K., X.Z., K.F., and M.G. did the experiments and analyzed the data. F.W. fabricated the microstructures and wrote the data analysis codes. DC and BM designed the simulations. P.S performed the simulations under the supervision of D.C. C.D. supervised the experimental work. B.M. supervised the theoretical work. F.W., D.C., B.M. & C.D wrote the paper.

## METHODS

### Experimental Procedure

#### On-chip experiments

Mask nanofabrication and PDMS microchamber patterning was done as described previously (Wu et al., 2015b). *E. coli* bacteria from a freezer stock were inoculated into M9 medium supplemented with 0.4% glycerol and 0.01% of protein hydrolysate amicase, and incubated overnight at 30 °C. The PDMS/glass chip was treated with oxygen plasma for 5 seconds to make the surface of the microchambers hydrophilic. 1 μl of the overnight bacterial culture was then pipetted onto the PDMS/glass chip that was clamped onto a custom-made baseplate. The droplet was then immediately covered by a 4.8% agarose pad supplemented with M9 broth, 0.4% glucose, 0.01% protein hydrolysate amicase, and 25 μg/ml cephalexin (Sigma-Aldrich). The baseplate was well sealed by a piece of parafilm to prevent drying and placed onto the microscope stage. For imaging the cell division event of *dnaC2(ts)/ΔslmA* cells, cephalexin was omitted in the agarose pad.

#### Fluorescence imaging

Widefield fluorescence imaging was carried out using Nikon Ti-E microscope with CFI Apo TIRF objective with an NA of 1.49. The microscope was enclosed by a custom-made chamber that was pre-heated overnight and kept at 39-40 °C. For excitation of mCerulean, sfGFP, mYPet, mCherry or mKate2 signal, cells were illuminated by Nikon-Intensilight illumination lamp through a CFP filter (λ_ex_ / λ_bs_ / λ_em_ =426-446 / 455 / 460-500 nm), YFP filter (λ_ex_ / λ_bs_ / λ_em_=490-510 / 515 / 520-550 nm), or an RFP filter cube (λ_ex_ / λ_bs_ / λ_em_ = 540-580 / 585 / 592 - 668). The fluorescence signal was recorded by an Andor iXon EMCCD camera. Images were acquired every 12 minutes for about 8 hours. The structured illumination images were taken using Nikon-Ti microscope equipped with a N-SIM module with a 100X objective (1.49), 515nm laser, and an Andor iXon EMCCD camera.

#### Image analysis and data analyses

Image analyses of wide-field images were carried out using our customized Matlab program, with automatic shape recognition and foci recognition. The data were plotted in Matlab, and if applicable, fitted with the curve fitting toolbox in Matlab. The Matlab code will be shared publicly upon publication of this paper.

#### Growth conditions

For genetic engineering, *E. coli* cells were incubated in Lysogeny broth (LB) supplemented, when required, with 100 μg/ml ampicillin (Sigma-Aldrich), 50 μg/ml kanamycin (Sigma-Aldrich), or 34 μg/ml chloramphenicol (Sigma-Aldrich) for plasmid selection, and with 25 μg/ml kanamycin or 11 μg/ml chloramphenicol for selection of the genomic insertions of gene cassettes. For on-chip experiments, we grew cells in liquid M9 minimum medium (Fluka Analytical) supplemented with 2 mM MgSO_4_, 0.1mM CaCl_2_, 0.4% glycerol (Sigma-Aldrich), and 0.01% protein hydrolysate amicase (PHA) (Fluka Analytical). For testing transertion, 34 μg/ml chloramphenicol (Sigma-Aldrich) and 100 μg/ml rifampicin are used in the agarose pad.

#### Strain construction

A list of strain is listed in the Supplemental Table 1. To construct FW2442, strain FW2177 was transduced with P1 phage JW5641 and selected for kanamycin resistance. The resulting strain was cured of kanamycin resistance by pCP20 and then transduced with P1 phage FW1957 and selected for kanamycin resistance and temperature sensitivity. To construct strain FW2502, strain FW2179 was transduced with P1 phage FW1363 and selected for chloramphenicol resistance. All the insertions were confirmed by sequencing.

## Theoretical Model

The 4.6 Mbp circular genome of *E. coli* was modeled as a polymer of beads that form a backbone chain consisting of a number (*n*_*b*_) of monomers, each with side-loop attached that contained n_s_ monomers, totaling 4 × 10^4^ beads (bead diameter σ = 0.04 μm, or 115 bp) that stretched out to a length of 1.6mm (Fig.2A). Here, each side loop was taken to be *n*_*s*_ = 62 beads long, which represented 7.2 kbp DNA that amounts to 2.4 μm of length that folded into a loop, and a main chain with *n*_*b*_ = 636 beads that corresponded to a length of 24.8 μm. Thus, the total chain length *l* = *n*_*b*_ σ + *n*_*b*_(*n*_*s*_ σ). The polymer was simulated by beads connected by a finitely extensible nonlinear elastic (FENE) potential

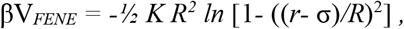

with *K* = *30* and *R* = *1.5* σ. The self-avoidance in the chain was incorporated via the repulsive part of the Lennard-Jones potential

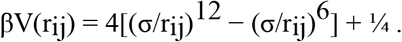

The presence of side loops generated both an effective bending stiffness as well as a “thickening” of the main chain. The latter effect led to a soft repulsion between spatially close but contour-wise distant parts of the chromosome. Both effects were well captured by approximating the soft effective repulsion between side-loops in terms of an excess Gaussian core (GC) potential
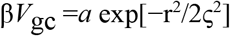

between the main-chain beads, in addition to the self-avoidance (Chaudhuri and Mulder, 2012). The interaction range between two side-loops is given by ς^2^ = 2 (*R*_*g*_)^2^ where *R*_*g*_ is the radius of gyration of side-loops given by *R*_*g*_= *c n*_*s*_^*3/5*^ σ where the numerical factor *c* = *0.323* was confirmed from independent molecular dynamics (MD) simulations (Chaudhuri and Mulder, 2012). Thus, the impact of side loops of length 2.4 μm can be incorporated through an additional GC interaction, of a width 0.21 μm and a strength proportional to the side-loop size, between backbone beads. Under strong confinement, in accordance with de Gennes’ blob picture, the interaction strength between loops is expected to grow with loop size (Jun and Mulder, PNAS 2006), and we assumed *a*= *n*_*s*_.

Crowders were modeled as non-additive depletants, so they did not interact amongst themselves but repel the beads of the polymer. To avoid introducing more interactions parameters, we assume this repulsion to be the same as that between monomers, having both repulsive Lennard-Jones and GC repulsion components.

The confinement is introduced through repulsive interaction between all beads (monomer and depletant) and walls of the confining cylindrical cell geometry. For this purpose, an integrated WCA repulsion
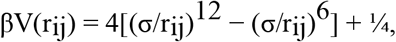

and Gaussian core with half the strength and width, *a/2* and ς/2 are used. To model a cylindrical cell of diameter 1μm, we used *D*=26.67σ and we varied the length of the cell.

To keep the density of depletants constant in a growing cell, we used a Widom insertion scheme that ensured that new depletants were added in a spatially homogeneously distributed manner near the simulated chromosome consistent with the equilibrium state. This was done by a trial move in a Monte-Carlo sense in which a new depletant particle was placed inside the cell within a 2ς range of the chain and the change in energy *ΔE* due to trial insertions was calculated. The insertion move was accepted with a probability proportional to the Boltzmann weight exp(-*βΔE*).

---

**Figure S1.**
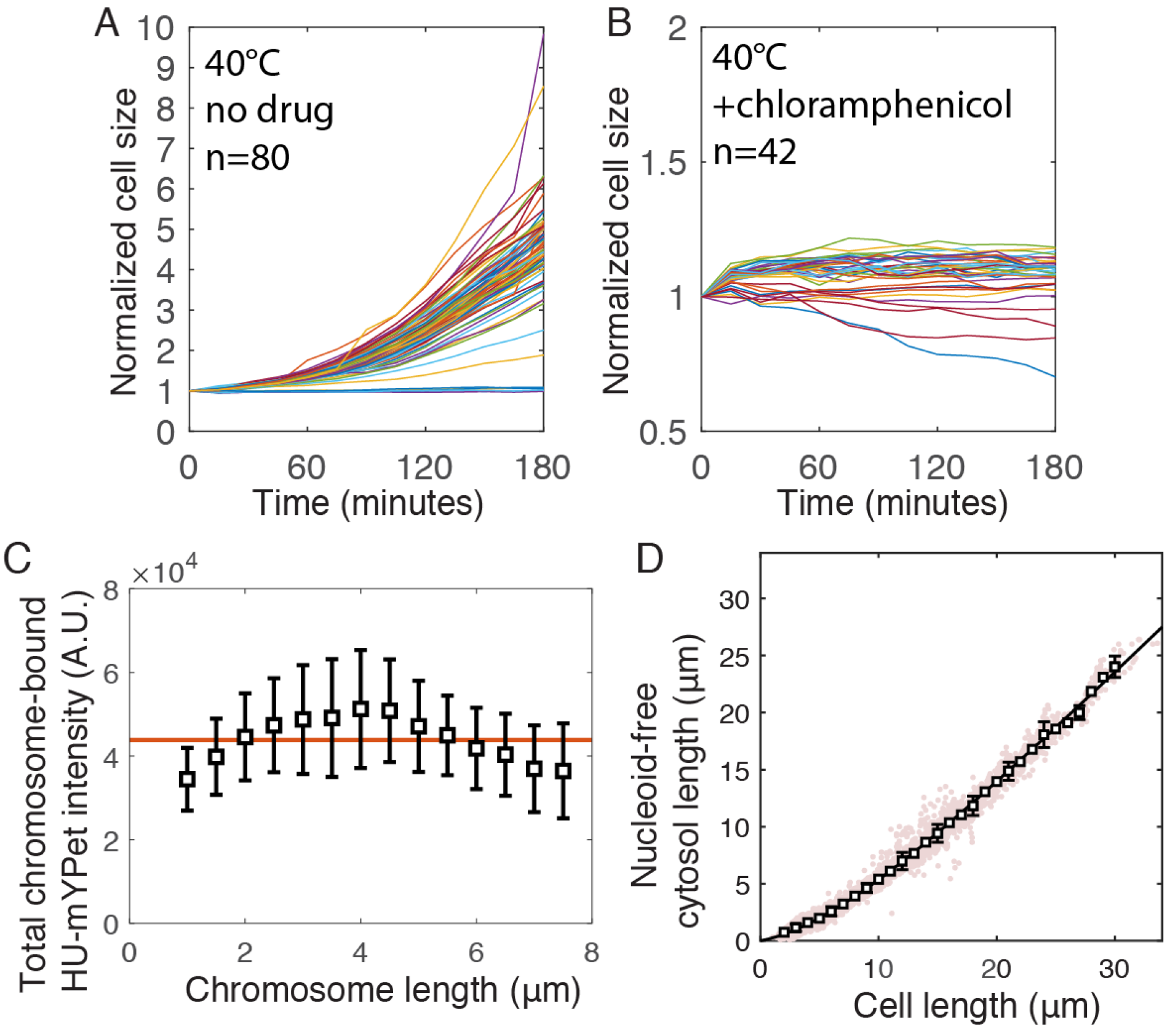
Cell growth with a single nucleoid requires protein synthesis and maintains a nucleoid-bound HU-mYPet level. A and B. Cell area measurement of *dnaC2(ts)* allel growing at non-permissive temperature without and with chloramphenicol treatment C. Cell length unoccupied by the single nucleoids in cylindrical *dnaC2(ts)* cells growing into different sizes. Error bars represent standard deviations. D. Total chromosome-bound HU-mYPet intensity in *dnaC2(ts)* cells growing into different sizes.

**Figure S2.**
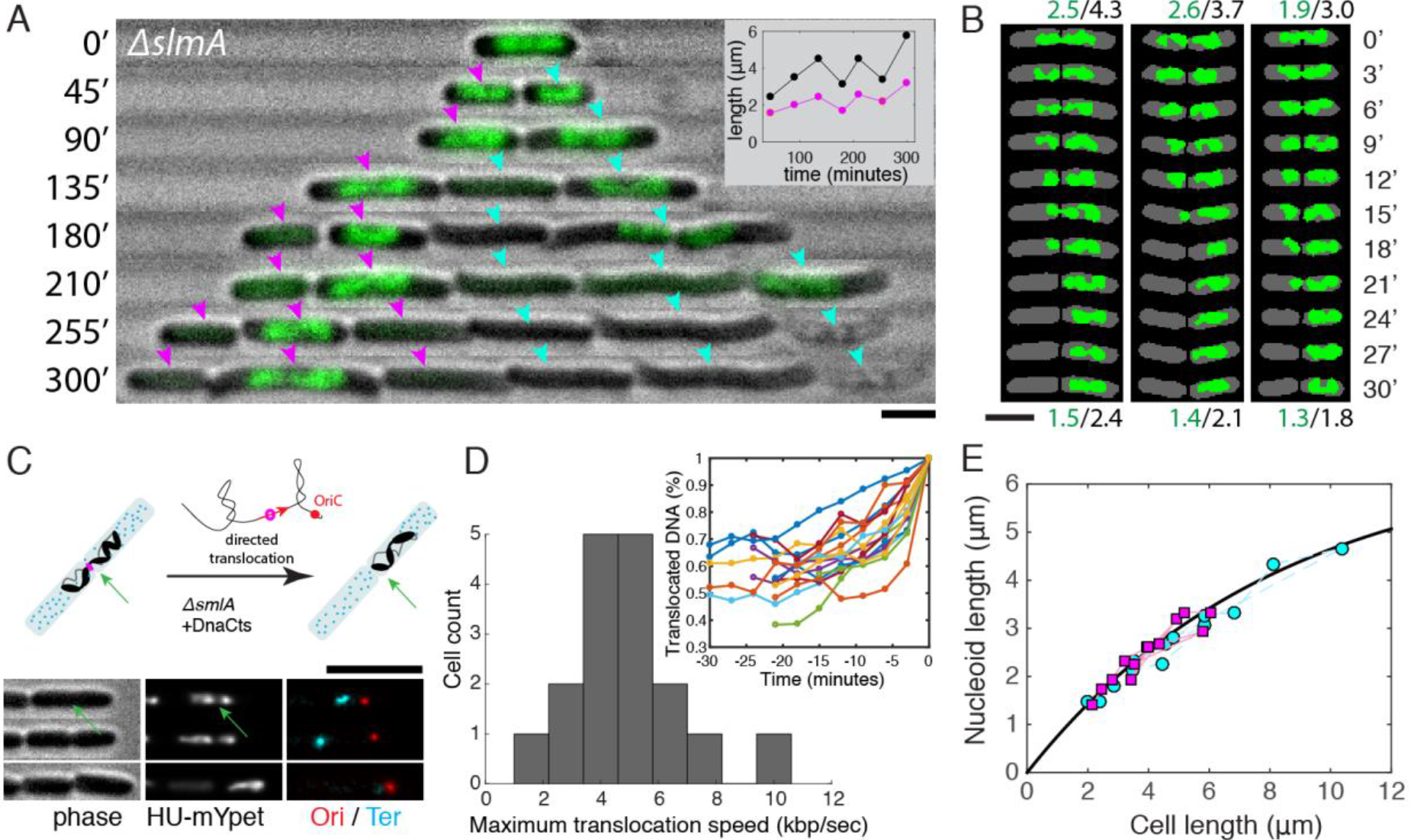
Cell division and chromosome contraction of *ΔslmA/dnaC2* mutant. A, time-lapse images nucleoid dynamics during the growth and division of *ΔslmA/dnaC2* cells growing at non-permissive temperature. Phase contrast in grey scale, and HU-mYPet in green. Time is labeled in minutes. Magenta and cyan arrows traces two cell lineages. Scale bars, 2 μm. Inset, cell size (black) and nucleoid size (magenta) change over time measured from the *nucleoid-containing* cell lineage at the left. B, three examples of chromosome translocation after cell constriction and prior to septation. Grey indicates automatically identified cell shape, and green indicates automatically identified nucleoid. C, illustration and time-lapse images showing that the chromosome translocation is oriented towards the cell half with the Ori focus. D, histogram showing the maximum DNA translocation speed estimated from the time-lapse fluorescent images. Inset, progression of DNA translocation in single cells over time. E, nucleoid/cell length relation in the two nucleoid-containing lineages indicated in A measured over two cell division events. Time interval is 15 minutes. Each had 13 time points. The black smooth line shows a section of the nucleoid-boundary response curve shown in Fig. 1C.

**Figure S3.**
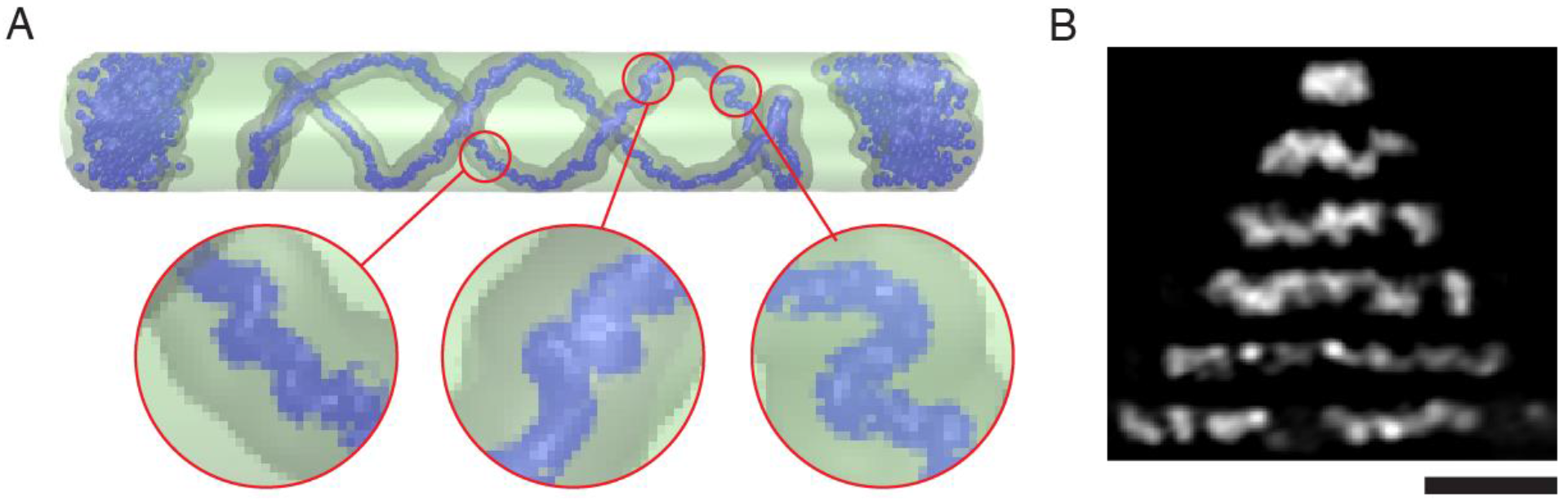
Effect of cell size on the nucleoid internal structure. A, snapshot of a model chromosome at cell length L=6 μm at a density of depletants of 212 μm^−3^, showing the polar segregation of the depletants and the helical backbone conformation. B, Structured Illumination Microscopy images of nucleoids of different lengths at their central focal planes. Scale bar, 2 μm.

**Figure S4.**
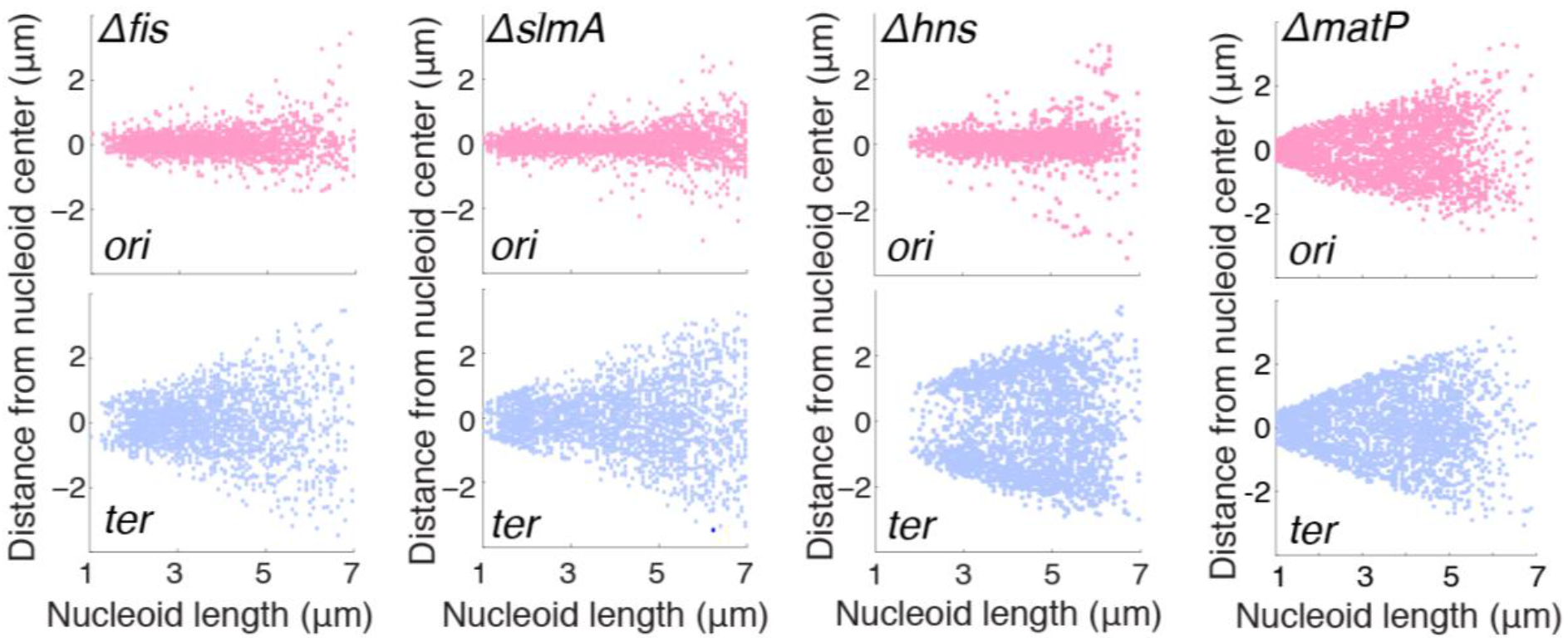
Ori/Ter foci positioning inside nucleoids of different lengths in different NAP mutants. Each panel displays the distances of Ori / Ter loci from the center of nucleoids as a function of the nucleoid length.

**Figure S5.**
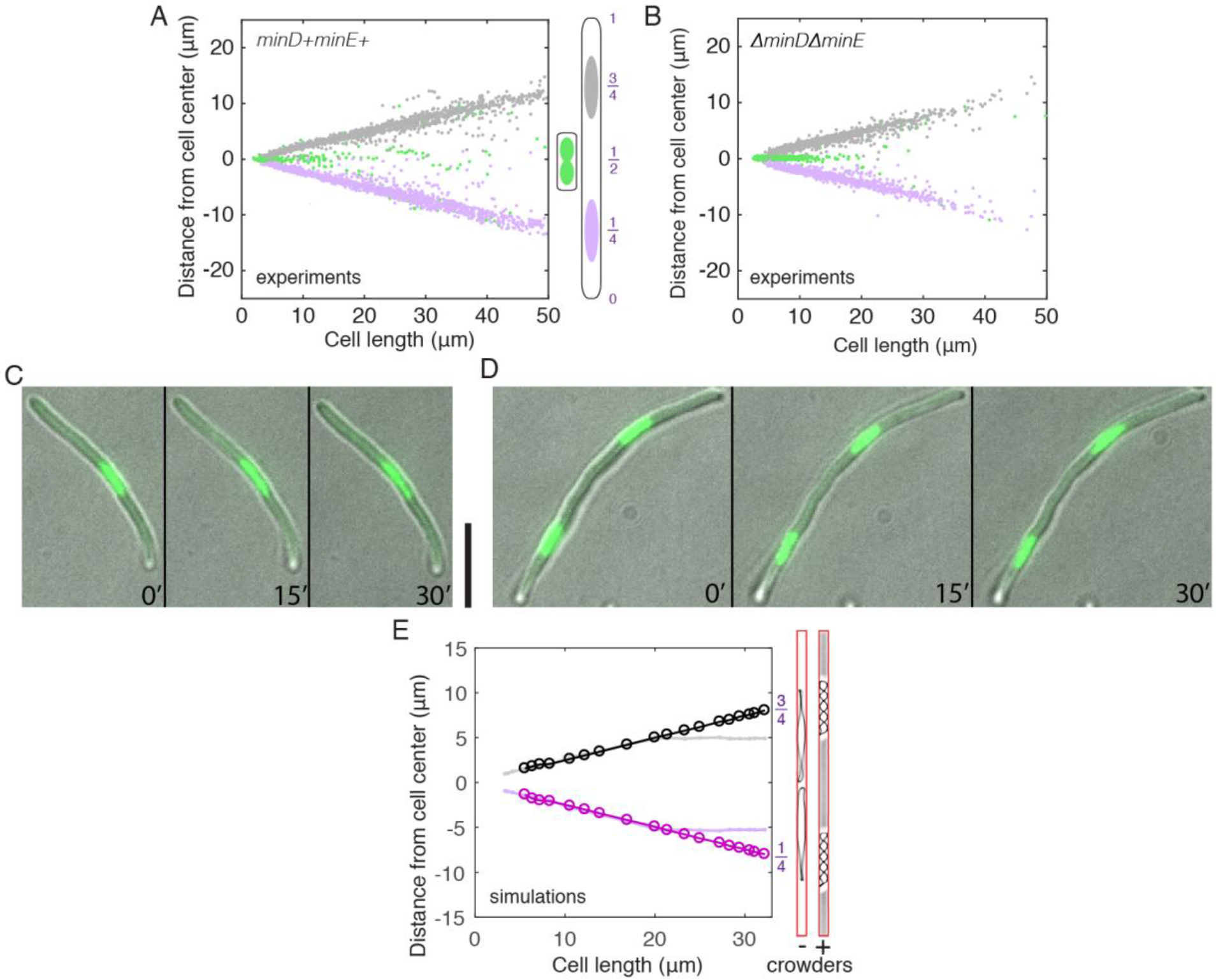
Sister chromosome positioning is not affected by abolishing transertion or Min proteins. A and B, distances of two sister chromosomes from the cell center in different cell lengths in *minDE*+ and *minDE*− cells (n = 3626). Green data points represent sister chromosomes that are still connected. Grey and purple data points indicate right and left chromosomes respectively. C and D, time-lapse images of single- or double-nucleoid cells treated by a combination of 34 μg/ml chloramphenicol and 100 μg/ml rifampicin, which were added into the agarose pad. Time 0’ is 10 minutes after inoculation onto the cover glass. Scale bar, 5 μm. E, distances of two sister chromosomes from cell center in different cell lengths obtained through simulations with (bright circles) and without (light lines) depletants.

**Supplementary Table T1.** List of strains used in this study.

| Strains | Descriptions | References |
| --- | --- | --- |
| JW5641 | BW25113, *ΔslmA::aph :: frt* | (Baba et al., 2006) |
| FW1957 | *dnaC2(ts) ΔmdoB::aph :: frt* | (Saifi and Ferat, 2012) |
| FW1363 | W3110, *ΔminDE::cat :: frt :: sacA* | (Wu et al., 2015b) |
| FW2177 | AB1157, *ori1:: lacOx240::hygR, ter3::tetOx240::accC1 ΔgalK::tetR-mCerulean :: frt, ΔleuB::lacI-mCherry :: frt, hupA-mYPet :: frt* | (Wu et al., 2018) |
| FW2179 | AB1157, *ori1:: lacOx240::hygR, ter3::tetOx240::accC1 ΔgalK::tetR-mCerulean :: frt, ΔleuB::lacI-mCherry :: frt, hupA-mYPet :: frt, dnaC2 (ts) :: aph frt* | (Wu et al., 2018) |
| FW2444 | AB1157, *ori1:: lacOx240::hygR, ter3::tetOx240::accC1 ΔgalK::tetR-mCerulean :: frt, ΔleuB::lacI-mCherry :: frt, hupA-mYPet :: frt, Δfis::frt, dnaC2 (ts) :: aph frt* | (Wu et al., 2018) |
| FW2479 | AB1157, *ori1:: lacOx240::hygR, ter3::tetOx240::accC1 ΔgalK::tetR-mCerulean :: frt, ΔleuB::lacI-mCherry :: frt, hupA-mYPet :: frt, Δhns::frt, dnaC2 (ts) :: aph frt* | (Wu et al., 2018) |
| FW2254 | AB1157, *ori1:: lacOx240::hygR, ter3::tetOx240::accC1 ΔgalK::tetR-mCerulean :: frt, ΔleuB::lacI-mCherry :: frt, hupA-mYPet :: frt, ΔmatP::frt, dnaC2 (ts) :: aph frt* | (Wu et al., 2018) |
| FW2442 | AB1157, *ori1:: lacOx240::hygR, ter3::tetOx240::accC1 ΔgalK::tetR-mCerulean :: frt, ΔleuB::lacI-mCherry :: frt, hupA-mYPet :: frt, ΔslmA::frt, dnaC2 (ts) :: aph frt* | This work |
| FW2502 | AB1157, *ori1:: lacOx240::hygR, ter3::tetOx240::accC1 ΔgalK::tetR-mCerulean :: frt, ΔleuB::lacI-mCherry :: frt, hupA-mYPet :: frt, ΔminDE::cat :: frt :: sacA, dnaC2 (ts) :: aph frt* | This work |

## References

Baba, T., Ara, T., Hasegawa, M., Takai, Y., Okumura, Y., Baba, M., Datsenko, K.A., Tomita, M., Wanner, B.L., and Mori, H. (2006). Construction of Escherichia coli K-12 in-frame, single-gene knockout mutants: the Keio collection. Mol Syst Biol 2, 2006.0008–2006.0008.

Badrinarayanan, A., Reyes-Lamothe, R., Uphoff, S., Leake, M.C., and Sherratt, D.J. (2012). In Vivo Architecture and Action of Bacterial Structural Maintenance of Chromosome Proteins. Science 338, 528.

Bernhardt, T.G., and de Boer, P.A.J. (2005). SlmA, a Nucleoid-Associated, FtsZ Binding Protein Required for Blocking Septal Ring Assembly over Chromosomes in E. coli. Mol Cell 18, 555–564.

Bickmore, Wendy A., and van Steensel, B. (2013). Genome Architecture: Domain Organization of Interphase Chromosomes. Cell 152, 1270–1284.

Bolzer, A., Kreth, G., Solovei, I., Koehler, D., Saracoglu, K., Fauth, C., Müller, S., Eils, R., Cremer, C., Speicher, M.R., et al. (2005). Three-Dimensional Maps of All Chromosomes in Human Male Fibroblast Nuclei and Prometaphase Rosettes. PLOS Biology 3, e157.

Cagliero, C., Grand, R.S., Jones, M.B., Jin, D.J., and O’Sullivan, J.M. (2013). Genome conformation capture reveals that the Escherichia coli chromosome is organized by replication and transcription. Nucleic Acids Res 41, 6058–6071.

Chaudhuri, D., and Mulder, B.M. (2012). Spontaneous Helicity of a Polymer with Side Loops Confined to a Cylinder. Physical Review Letters 108, 268305.

Dame, R.T., Noom, M.C., and Wuite, G.J.L. (2006). Bacterial chromatin organization by H-NS protein unravelled using dual DNA manipulation. Nature 444, 387–390.

Danilova, O., Reyes-Lamothe, R., Pinskaya, M., Sherratt, D., and Possoz, C. (2007). MukB colocalizes with the oriC region and is required for organization of the two Escherichia coli chromosome arms into separate cell halves. Mol Microbiol 65, 1485–1492.

de Vries, R. (2010). DNA condensation in bacteria: interplay between macromolecular crowding and nucleoid proteins. Biochimie 92, 1715–1721.

Di Ventura, B., Knecht, B., Andreas, H., Godinez, W.J., Fritsche, M., Rohr, K., Nickel, W., Heermann, D.W., and Sourjik, V. (2013). Chromosome segregation by the Escherichia coli Min system. Mol Syst Biol 9.

Dupaigne, P., Tonthat, N.K., Espéli, O., Whitfill, T., Boccard, F., and Schumacher, M.A. (2012). Molecular basis for a protein-mediated DNA-bridging mechanism that functions in condensation of the E. coli chromosome. Mol Cell 48, 560–571.

Ellis, R.J. (2001). Macromolecular crowding: an important but neglected aspect of the intracellular environment. Current Opinion in Structural Biology 11, 114–119.

Espéli, O., Borne, R., Dupaigne, P., Thiel, A., Gigant, E., Mercier, R., and Boccard, F. (2012). A MatP-divisome interaction coordinates chromosome segregation with cell division in E. coli. EMBO J 31, 3198–3211.

Fisher, J.K., Bourniquel, A., Witz, G., Weiner, B., Prentiss, M., and Kleckner, N. (2013). FourDimensional Imaging of E. coli Nucleoid Organization and Dynamics in Living Cells. Cell 153, 882–895.

Ganji, M., Shaltiel, I.A., Bisht, S., Kim, E., Kalichava, A., Haering, C.H., and Dekker, C. (2018). Realtime imaging of DNA loop extrusion by condensin. Science.

Hussain, S., Wivagg, C.N., Szwedziak, P., Wong, F., Schaefer, K., Izoré, T., Renner, L.D., Holmes, M.J., Sun, Y., Bisson-Filho, A.W., et al. (2018). MreB filaments align along greatest principal membrane curvature to orient cell wall synthesis. eLife 7, e32471.

Jun, S., and Mulder, B. (2006). Entropy-driven spatial organization of highly confined polymers: Lessons for the bacterial chromosome. Proc Natl Acad Sci USA 103, 12388–12393.

Kellenberger, E., Ryter, A., and Sechaud, J. (1958). Electron microscope study of DNA-containing plasms. II. Vegetative and mature phage DNA as compared with normal bacterial nucleoids in different physiological states. J Biophys Biochem Cytol 4, 671–678.

Kuwada, N.J., Cheveralls, K.C., Traxler, B., and Wiggins, P.A. (2013). Mapping the driving forces of chromosome structure and segregation in Escherichia coli. Nucleic Acids Res 41, 7370–7377.

Lesterlin, C., Ball, G., Schermelleh, L., and Sherratt, D.J. (2013). RecA bundles mediate homology pairing between distant sisters during DNA break repair. Nature 506, 249.

Lioy, V.S., Cournac, A., Marbouty, M., Duigou, S., Mozziconacci, J., Espéli, O., Boccard, F., and Koszul, R. (2018). Multiscale Structuring of the E. coli Chromosome by Nucleoid-Associated and Condensin Proteins. Cell.

Luijsterburg, M.S., Noom, M.C., Wuite, G.J.L., and Dame, R.T. (2006). The architectural role of nucleoid-associated proteins in the organization of bacterial chromatin: A molecular perspective. J Struct Biol 156, 262–272.

Männik, J., Bailey, M.W., O’Neill, J.C., and Männik, J. (2017). Kinetics of large-scale chromosomal movement during asymmetric cell division in Escherichia coli. PLoS Genet 13, e1006638.

Mercier, R., Petit, M., Schbath, S., Robin, S., Karoui, M.E., Boccard, F., and Espéli, O. (2008). The MatP/matS site-specific system organizes the terminus region of the E. coli chromosome into a macrodomain. Cell 135, 475–485.

Minc, N., Burgess, D., and Chang, F. (2011). Influence of Cell Geometry on Division-Plane Positioning. Cell 144, 414–426.

Niki, H., Yamaichi, Y., and Hiraga, S. (2000). Dynamic organization of chromosomal DNA in Escherichia coli. Genes Dev 14, 212–223.

Nolivos, S., Upton, A.L., Badrinarayanan, A., Müller, J., Zawadzka, K., Wiktor, J., Gill, A., Arciszewska, L., Nicolas, E., and Sherratt, D. (2016). MatP regulates the coordinated action of topoisomerase IV and MukBEF in chromosome segregation. Nat Commun 7, 10466.

Pan, C.Q., Finkel, S.E., Cramton, S.E., Feng, J.-A., Sigman, D.S., and Johnson, R.C. (1996). Variable Structures of Fis-DNA Complexes Determined by Flanking DNA – Protein Contacts. Journal of Molecular Biology 264, 675–695.

Parry, Bradley R., Surovtsev, Ivan V., Cabeen, Matthew T., O’Hern, Corey S., Dufresne, Eric R., and Jacobs-Wagner, C. (2014). The Bacterial Cytoplasm Has Glass-like Properties and Is Fluidized by Metabolic Activity. Cell 156, 183–194.

Peeters, E., Driessen, R.P.C., Werner, F., and Dame, R.T. (2015). The interplay between nucleoid organization and transcription in archaeal genomes. Nature Reviews Microbiology 13, 333.

Pelletier, J., Halvorsen, K., Ha, B.-Y., Paparcone, R., Sandler, S.J., Woldringh, C.L., Wong, W.P., and Jun, S. (2012). Physical manipulation of the Escherichia coli chromosome reveals its soft nature. Proc Natl Acad Sci USA 109, E2649–E2656.

Postow, L., Hardy, C.D., Arsuaga, J., and Cozzarelli, N.R. (2004). Topological domain structure of the Escherichia coli chromosome. Genes Dev 18, 1766–1779.

Rathgeber, S., Pakula, T., Wilk, A., Matyjaszewski, K., and Beers, K.L. (2005). On the shape of bottlebrush macromolecules: Systematic variation of architectural parameters. J Chem Phys 122.

Reiss, P., Fritsche, M., and Heermann, D.W. (2011). Looped star polymers show conformational transition from spherical to flat toroidal shapes. Phys Rev E 84, 051910.

Reyes-Lamothe, R., Possoz, C., Danilova, O., and Sherratt, D.J. (2008). Independent Positioning and Action of Escherichia coli Replisomes in Live Cells. Cell 133, 90–102.

Saifi, B., and Ferat, J.-L. (2012). Replication Fork Reactivation in a dnaC2 Mutant at Non-Permissive Temperature in Escherichia coli. PLoS ONE 7, e33613.

Saleh, O.A., Pérals, C., Barre, F.-X., and Allemand, J.-F. (2004). Fast, DNA-sequence independent translocation by FtsK in a single-molecule experiment. EMBO J 23, 2430–2439.

Schneider, R., Travers, A., and Muskhelishvili, G. (1997). FIS modulates growth phase-dependent topological transitions of DNA in Escherichia coli. Mol Microbiol 26, 519–530.

Si, F., Li, D., Cox, S.E., Sauls, J.T., Azizi, O., Sou, C., Schwartz, A.B., Erickstad, M.J., Jun, Y., Li, X., et al. (2017). Invariance of Initiation Mass and Predictability of Cell Size in Escherichia coli. Current Biology 27, 1278–1287.

Stillinger, F.H. (1976). Phase transitions in the Gaussian core system. The Journal of Chemical Physics 65, 3968–3974.

Umbarger, Mark A., Toro, E., Wright, Matthew A., Porreca, Gregory J., Baù, D., Hong, S.-H., Fero, Michael J., Zhu, Lihua J., Marti-Renom, Marc A., McAdams, Harley H., et al. (2011). The ThreeDimensional Architecture of a Bacterial Genome and Its Alteration by Genetic Perturbation. Mol Cell 44, 252–264.

van Loenhout, M.T.J., de Grunt, M.V., and Dekker, C. (2012). Dynamics of DNA Supercoils. Science 338, 94–97.

van Noort, J., Verbrugge, S., Goosen, N., Dekker, C., and Dame, R.T. (2004). Dual architectural roles of HU: formation of flexible hinges and rigid filaments. Proc Natl Acad Sci U S A 101, 6969–6974.

Wang, S., Moffitt, J.R., Dempsey, G.T., Xie, X.S., and Zhuang, X. (2014). Characterization and development of photoactivatable fluorescent proteins for single-molecule-based superresolution imaging. Proc Natl Acad Sci USA 111, 8452–8457.

Wang, X., Liu, X., Possoz, C., and Sherratt, D.J. (2006). The two Escherichia coli chromosome arms locate to separate cell halves. Genes Dev 20, 1727–1731.

Weber, S.C., Spakowitz, A.J., and Theriot, J.A. (2010). Bacterial Chromosomal Loci Move Subdiffusively through a Viscoelastic Cytoplasm. Physical Review Letters 104, 238102.

Wery, M., Woldringh, C.L., and Rouviere-Yaniv, J. (2001). HU-GFP and DAPI co-localize on the Escherichia coli nucleoid. Biochimie 83, 193–200.

Wiggins, P.A., Cheveralls, K.C., Martin, J.S., Lintner, R., and Kondev, J. (2010). Strong intranucleoid interactions organize the Escherichia coli chromosome into a nucleoid filament. Proc Natl Acad Sci USA 107, 4991–4995.

Woldringh, C.L. (2002). The role of co-transcriptional translation and protein translocation (transertion) in bacterial chromosome segregation. Mol Microbiol 45, 17–29.

Woldringh, C.L., Hansen, F.G., Vischer, N.O.E., and Atlung, T. (2015). Segregation of chromosome arms in growing and non-growing Escherichia coli cells. Front Microbiol 6.

Wu, F., Japaridze, A., Zheng, X., Kerssemakers, J.W.J., and Dekker, C. (2018). Direct Imaging of the circular chromosome of a live bacterium. bioRxiv.

Wu, F., van Rijn, E., van Schie, B.G.C., Keymer, J.E., and Dekker, C. (2015a). Multicolor imaging of bacterial nucleoid and division proteins with blue, orange and near-infrared fluorescent proteins. Front Microbiol 6, 607.

Wu, F., van Schie, B.G.C., Keymer, J.E., and Dekker, C. (2015b). Symmetry and scale orient Min protein patterns in shaped bacterial sculptures. Nat Nanotechnol 10, 719–726.

Wu, K.-T., Hishamunda, J.B., Chen, D.T.N., DeCamp, S.J., Chang, Y.-W., Fernández-Nieves, A., Fraden, S., and Dogic, Z. (2017). Transition from turbulent to coherent flows in confined three-dimensional active fluids. Science 355.

Yamaichi, Y., and Niki, H. (2004). migS, a cis-acting site that affects bipolar positioning of oriC on the Escherichia coli chromosome. EMBO J 23, 221–233.

Young, K.D. (2006). The selective value of bacterial shape. Microbiology and Molecular Biology Reviews 70, 660–703.

Youngren, B., Nielsen, H.J., Jun, S., and Austin, S. (2014). The multifork Escherichia coli chromosome is a self-duplicating and self-segregating thermodynamic ring polymer. Genes Dev 28, 71–84.

Zhou, H.-X., Rivas, G., and Minton, A.P. (2008). Macromolecular crowding and confinement: biochemical, biophysical, and potential physiological consequences. Annual review of biophysics 37, 375–397.

